# Mechanisms of Antimicrobial Agent Cetylpyridinium Chloride Mitochondrial Toxicity in Rodent and Primary Human Cells: Super-resolution Microscopy Reveals Nanostructural Disruption

**DOI:** 10.1101/2022.09.27.509813

**Authors:** Sasha R. Weller, John E. Burnell, Brandon M. Aho, Bright Obeng, Emily L. Ledue, Juyoung K. Shim, Samuel T. Hess, Julie A. Gosse

**Affiliations:** Department of Molecular and Biomedical Sciences, University of Maine, Orono, ME, USA; Department of Physics and Astronomy, University of Maine, Orono, ME, USA; Department of Biology, University of Maine Augusta, Augusta, ME, USA

**Author notes:** Corresponding authors: Julie A. Gosse, Samuel T. Hess. These authors contributed equally (Sasha R. Weller, John E. Burnell, Brandon M. Aho).

**Keywords:** cetylpyridinium chloride, antibacterial agent, mitochondria, super-resolution microscopy, keratinocyte, NIH-3T3, mast cell, RBL-2H3, mitochondrial Ca^2+^, ATP, oxygen consumption rate, fluorescence photoactivation localization microscopy (FPALM), mitotoxicity

## Abstract

People are exposed to high concentrations of antibacterial agent cetylpyridinium chloride (CPC) via personal care and food products, despite little information regarding CPC effects on eukaryotes. CPC is used as an antibacterial agent via a detergent mechanism when above ∼600- 900 μM. While three previous studies suggested CPC mitochondrial toxicity, this phenomenon is not well-studied. Here, we show that low-micromolar CPC inhibits mitochondrial ATP production in primary human keratinocytes, mouse NIH-3T3 fibroblasts, and rat RBL-2H3 immune mast cells, in galactose media, which causes cells to produce ATP via mitochondria. ATP inhibition via CPC (EC_50_ 1.7 µM) is nearly as potent as that caused by canonical mitotoxicant CCCP (EC_50_ 1.2 µM). CPC inhibition of oxygen consumption rate (OCR) tracks with that of ATP: OCR is halved due to 1.75 μM CPC in RBL-2H3 cells and 1.25 μM in primary human keratinocytes. Here we demonstrate that CPC is more potent as a mitotoxicant than as an immune mast cell signaling inhibitor, an effect published previously. Mitochondrial [Ca^2+^] changes can cause mitochondrial dysfunction. Here we show, using a novel plate reader assay with reporter CEPIA2mt, that CPC causes mitochondrial Ca^2+^ efflux from mast cells via an ATP-inhibition mechanism. Using super-resolution microscopy (fluorescence photoactivation localization) in live cells, we have discovered that CPC causes mitochondrial nanostructural defects in fibroblasts, including the formation of spherical structures with donut-like cross section, as quantified by novel Fourier transform analysis. This work reveals CPC as a mitotoxicant despite widespread use, highlighting the importance of further research into its toxicological safety.

## Introduction

Cetylpyridinium chloride (CPC) is a quaternary ammonium compound (QAC) and antimicrobial used in personal care and food products at a concentration of ∼1,500 μM - 3,000 μM (Rawlinson *et al*., 2008) since the 1930s (ACS, 2021). CPC is also used directly on human food in agricultural processing (HHS, 2004) and in cleaning products. For example, CPC lowers *Salmonella* and *Listeria* levels on meat (İlhak *et al*., 2018) and vegetables (Wang *et al*., 2001). Thus, human and environmental exposure must occur; however, little is known about human CPC body burden.

CPC is likely bioavailable (Van Leeuwen *et al*., 2015) and absorbable via the skin and gastrointestinal tract (Voutchkova *et al*., 2010). Its LogP_octanol/water_ partition ratio (PubChem) suggests significant oral bioavailability (Klaassen and Casarett, 2019), though no publications to date have examined CPC levels in human tissues or blood. A study in rats found 0.26 μM CPC in blood shortly after 1 mg/kg oral administration (Pottel *et al*., 2020) and 0.1 μM after 20 h. There exists evidence that CPC is retained in oral mucosa of humans who rinsed with and expelled 10 mL of 2200 μM CPC mouthwash for 1 min: saliva still contained 100 μM CPC after 30 min, and 1 μM even 24 h later (Bonesvoll and Gjermo, 1978). In the current study, we exposed cells to such CPC doses (low-μM). Similar QAC compounds are detected in the blood of 80% of humans sampled though CPC was not measured (Hrubec *et al*., 2021). Indeed, CPC is understudied compared to other QACs (based upon thorough PubMed searches).

Further CPC exposure estimates can be made by focusing on chicken consumption due to the presence of CPC in chicken processing (HHS, 2004; Yang and Chuan, 2016). A typical food-processing procedure involving a 1 min submergence of chicken in a 0.0125% CPC bath led to an average residual level of 11.7 mg/kg CPC in the chicken (Zhou *et al*., 1999). The CPC solution limit for agricultural processing is 8% concentration by weight (FDA, 2020a); extrapolating to this limit, we calculate that chicken meat treated maximally would retain 749 mg/kg CPC. The average American person consumes 0.12 kg of chicken per day (Council, 2022), which would result in 89.9 mg of CPC consumed daily if all their chicken had been CPC-treated. An industry survey indicated that roughly 10% of U.S. chicken is treated with CPC (McKee, 2011). Assuming an average American adult weight of 84 kg (CDC, 2021) and if all consumed chicken were from of the 10% treated with CPC, this daily chicken consumption would result in an exposure of ∼1 mg CPC/kg body weight. In a mouse study, an oral dosage of 1 mg/kg led to a peak concentration of 0.26 μM CPC in mouse blood and ∼0.1 μM 24 hr later (Pottel *et al*., 2020). Based on these calculations, we could expect a typical American to have a blood CPC concentration ranging from ∼0.1 μM - 0.3 μM due to chicken consumption alone, which is within the range of the 0.01 μM - 0.15 μM of other QAC varieties that have been detected in human blood (Hrubec *et al*., 2021). Regarding potential environmental exposures to humans and wildlife, CPC has been detected at levels up to 0.15 μM in rivers and municipal wastewater (Shrivas and Wu, 2007). Furthermore, related QACs have also been found in estuary sediment (Li and Brownawell, 2010) and in human sewage (Ruan *et al*., 2014). These emerging data points and considerations highlight the importance of research to determine human and wildlife exposure levels to CPC.

Classically, CPC has been employed for antibacterial applications in over-the-counter medicinal products (PubChem) like anti-gingival mouthwashes, toothpastes, lozenges, lip balms, and others (Mao *et al*., 2020). CPC helps to fight plaque (Sreenivasan *et al*., 2013), gingivitis (Teng *et al*., 2016), halitosis (Yaegaki and Sanada, 1992), and bacterial infections (Schaeffer *et al*., 2011; Latimer *et al*., 2015). Due to these oral care properties, including potential effects on bone signaling (Zheng *et al*., 2013), the addition of CPC to dental restorations is also being explored (Tomino *et al*., 2016; Yamamoto *et al*., 2022). CPC is used in numerous other cosmetics (hair coloring and styling products, deodorant, hair conditioners, and cosmetic biocides) and in the active pharmaceutical industry as an intermediate (PubChem). An unfortunate downside of CPC usage as an antibacterial could potentially be the development of antibiotic-resistant bacterial strains, such as those found following exposure to other (non-CPC) QACs (Soumet *et al*., 2016), including resistance to the antibiotic ciprofloxacin in *Listeria* (Guerin *et al*., 2021) and *E. coli* (Maertens *et al*., 2020). Furthermore, of the clinical isolates from patients infected with *E. coli*, those with greater resistance to QACs had greater resistance to antibiotics (Buffet-Bataillon *et al*., 2011). Thus, the use of CPC as an antibacterial may exacerbate the development of antibiotic resistant strains of bacteria, though CPC-specific studies of antibacterial resistance are limited.

The antibacterial properties of CPC stem from both its positively-charged headgroup and its hydrophobic tail: CPC acts as a detergent that lyses bacteria when above its critical micelle concentration of ∼600 μM - 900 μM (Mandal and Nair, 2002; Varade *et al*., 2005; Abezgauz *et al*., 2010; Shi *et al*., 2011; Shahinas *et al*., 2015).

Newer potential positive uses for CPC seem to be emerging: antiparasitic and antiviral. CPC executes antiparasitic actions against the nematode parasite *Strongyloides ratti* (10-100 μM exposures) (Keiser and Häberli, 2021), as well as against the amoeba parasite *Entamoeba histolytica* (25 μM exposure) (Shahinas *et al*., 2015). CPC displays antiviral activity through direct interaction with virus particles (Popkin *et al*., 2017; Seo *et al*., 2019; Koch-Heier *et al*., 2021; Bañó-Polo *et al*., 2022). For example, direct CPC exposure is virucidal to influenza viruses *in vitro* (EC_50_: 15-37 μM; exposure: 5 min) (Popkin *et al*., 2017). Moreover, mortality of influenza-infected mice (Popkin *et al*., 2017) and of zebrafish (0.1 μM for 1 hr) was reduced by CPC exposure (Raut *et al*., 2022). This suggests that CPC has potential use as an antiviral agent against influenza.

Further antiviral CPC actions are known or under current investigation. A small-scale human study indicated that CPC may treat upper respiratory infections (Mukherjee *et al*., 2017). Furthermore, there are currently several human clinical trials which assess the efficacy of CPC against SARS-CoV2 (NIH, 2022). The ongoing, deadly SARS-CoV2 pandemic (Organization, 2020) highlights the necessity of effective antivirals. For example, CPC in mouthwash (Muñoz-Basagoiti *et al*., 2020) and in lozenge form (Steyer *et al*., 2021) is effective against SARS-CoV2 (Alemany *et al*., 2022): CPC lyses viral particles (Koch-Heier *et al*., 2021; Bañó-Polo *et al*., 2022) and inhibits viral fusion (Muñoz-Basagoiti *et al*., 2020). Importantly, CPC’s virucidal potency is not dampened in the presence of human saliva nor by current viral variants (Muñoz-Basagoiti *et al*., 2020; Anderson *et al*., 2022).

While these beneficial antibacterial and antiviral effects are well-documented, CPC does lack antibacterial efficacy compared to plain cleansers, in hand soap and hand sanitizer formulations; this insufficiency led the FDA to effectively ban its presence in these products in 2016 (Wolf, 2016) and 2019 (Gottlieb, 2019), respectively. However, as noted above, CPC is still permitted as a food additive in chicken processing (FDA, 2020b) and in numerous personal care and household products (PubChem). Despite a continuing range of uses, little is known about CPC effects on eukaryotes. Thus, there is a growing need for studies on CPC eukaryotic toxicity.

Despite a lack of data on potential adverse human health effects of CPC exposure, there are a few known eukaryotic effects (Burnell, 2022). Intriguingly, QAC compounds have been shown to be toxic to mitochondria in living humans: extracted white blood cells exhibited decreased oxygen consumption rates with increasing QAC exposure levels (Hrubec *et al*., 2021). However, the Hrubec study did not examine CPC specifically. CPC structure alone implicates eukaryotic toxicity because its 16-carbon tail is the length found to be maximally toxic in QACs compared to other chain lengths (Moss and Mayo-Bean, 2010). Additionally, CPC is a lipophilic cation, a type of chemical known to be targeted to and to accumulate in mitochondria (Murphy, 2008).

Mitochondria are most known for their role in the production of ATP through oxidative phosphorylation in eukaryotic cells. This process is dependent on the functioning of mitochondrial complexes I-IV and the lipid ubiquinone (the electron transport chain, ETC). Complexes I, III, and IV use Gibbs free energy derived from the transfer of electrons from reduced electron carriers through the chain to terminal electron acceptor oxygen (respiration) in order to actively pump protons against their concentration gradient from the mitochondrial matrix into the intermembrane space. These protons flow down their electrochemical gradient back into the matrix through ATP synthase, which drives the formation of ATP from ADP and inorganic phosphate. Thus, oxygen consumption rate (OCR) can be measured to assess chemical effects on mitochondrial function.

One hallmark of cancer cells is the “Warburg effect,” which is the observation that these cells increase ATP production through the mitochondrial-independent process of glycolysis rather than through oxidative phosphorylation (Nelson and Cox, 2017). Despite the Warburg effect preference for glycolytic ATP production, immortalized cell cultures can rely on oxidative phosphorylation for ATP production (Rossignol *et al*., 2004). The presence or absence of glucose can be used to assess reliance on the mitochondria to produce ATP, with the substitution of glucose in cell media with its epimer galactose exploiting this difference.

Galactose metabolism can feed into the glycolytic pathway: galactose is converted to glucose 1-phosphate, via the Leloir Pathway (Holden *et al*., 2003), which is fed into the glycolytic pathway by glucose 1-phosphate conversion to glucose 6-phosphate by phosphoglucomutase (Nelson and Cox, 2017). However, eukaryotes preferentially use galactose in biosynthetic pathways (Reitzer *et al*., 1979), rather than in glycolysis, when amino acids are present. This avoidance of glycolysis when galactose is the only sugar present occurs because ATP production via mitochondria can be sufficiently and preferentially covered by amino acid metabolism via the TCA cycle and mitochondrial respiration (Deberardinis *et al*., 2008; Dang, 2010). For example, the amino acid glutamine can provide a majority of HeLa cells’ needed ATP (Reitzer *et al*., 1979). This preference for amino acids over galactose for ATP production may stem from the faster catabolism of amino acids compared to that of galactose (Lowry and Passonneau, 1969; Mulhausen and Mendicino, 1970). Thus, media containing galactose and glutamine is used experimentally to interrogate agents’ effects on mitochondrial ATP production.

Mitochondria play roles beyond that of metabolism. In healthy cells, mitochondria are involved in cell death and in cellular Ca^2+^ signaling dynamics (Jeong and Seol, 2008), which are essential for the functioning of the immune (Vig and Kinet, 2009) and nervous systems (Kawamoto *et al*., 2012). Mitochondrial Ca^2+^ levels are crucial in mitochondrial function. For example, mitochondrial matrix Ca^2+^ activates a cohort of metabolic enzymes, including glycerol- 3-phosphate dehydrogenase (Wernette *et al*., 1981) involved in lipid (Zheng *et al*., 2019) and other metabolism (Mráček *et al*., 2013), isocitrate dehydrogenase (Denton *et al*., 1978) of the TCA cycle, and regulatory enzymes (Kolobova *et al*., 2001; Trias *et al*., 2017) such as pyruvate dehydrogenase phosphatase (Denton *et al*., 1972). Additionally, protein S100A1, present in the brain, cardiac, and skeletal muscle tissues (Engelkamp *et al*., 1992), binds to ATP synthase and elevates ATP production in a mitochondrial Ca^2+^-dependent manner (Boerries *et al*., 2007). Altogether, these findings implicate mitochondrial Ca^2+^ in the regulation of metabolism.

In the current study, mitochondrial Ca^2+^ was assayed with reporter protein construct CEPIA2mt rather than with organic dyes. Fluorescent reporter construct proteins possess structure which protects the internal fluorophore from photophysical interference by exogenous agents (Brejc *et al*., 1997; Weatherly *et al*., 2016; Weatherly *et al*., 2018)

Normal mitochondrial nanostructure, typically involving networks of fused mitochondria, is essential for healthy cell function. Mitochondrial Ca^2+^ dynamics, ATP production, and cellular respiration are maintained in part by cycles of fusion and fission. Mitochondria undergo fission or fusion in response to metabolic changes arising from environmental or toxic stressors: fusion to share mitochondrial components such as proteins or lipids, and fission to meet organelle demand in a growing cell or to clear dysfunctional mitochondria (Youle and Van Der Bliek, 2012). Mitochondrial uncouplers such as carbonyl cyanide 3-chlorophenylhydrazone (CCCP) and carbonyl cyanide *p*-triflouromethoxyphenylhydrazone (FCCP) increase fission (Griparic *et al*., 2007; Giedt *et al*., 2012). Uncouplers induce mitochondrial swelling (Hanstein, 1976) and formation of donut-shaped mitochondria when followed by microtubule release and aberrant fusion (Liu and Hajnoczky, 2011). To assess whether CPC impacts mitochondrial ultrastructure, this study uses the live-cell super-resolution method fluorescence photoactivation localization microscopy (FPALM). Environmentally-induced mitochondrial dysfunction can lead to various health problems, including cardiac disease (Schwarz *et al*., 2014), obesity (Heinonen *et al*., 2015), and Alzheimer’s disease (Yan *et al*., 2013), to name a few.

To date, there are three initial studies which indicate CPC mitochondrial toxicity (Saladino *et al*., 1971; Chávez and Concepcion, 1982; Datta *et al*., 2017b). The first utilized isolated rat mitochondria and found that CPC reduced OCR, possibly via direct complex I inhibition and collapse of the mitochondrial membrane potential (MMP) (Chávez and Concepcion, 1982). Their use of isolated mitochondria as a model, while important for defining mitotoxicants, is a limitation because isolated mitochondria are known to exaggerate toxic effects due to their fragility (Picard *et al*., 2011). In addition, Chávez *et al*. used high CPC doses, thus confounding interpretation of whether relevant, non-cytotoxic exposures to CPC are mitotoxic.

A second study used whole cells rather than isolated mitochondria: human Leber’s hereditary optic neuropathy (LHON) osteosarcoma cytoplasmic hybrid cells (cybrids) (Datta *et al*., 2017b). LHON cells contain a mutation which may exaggerate test chemical toxicity. LHON is a disease of progressive blindness due to cell death caused by mutations of mitochondrial complex I (Kirches, 2011). LHON mutations are known to affect the performance of mitochondrial complex I in the ability of mitochondria to produce ATP (Baracca *et al*., 2005). Supporting this conclusion, a separate study by the same authors revealed LHON cybrids to be more sensitive to the mitotoxic effect of a related QAC, benzalkonium chloride, compared to cells lacking the mutation (Datta *et al*., 2017a). Thus, the use of LHON cells may risk exaggeration of CPC mitoxicity, compared to the chemical’s effects in normal cells. In this cell type, CPC decreased ATP production following a 24 hr exposure (Datta *et al*., 2017b); the accompanying cell death measurement was qualitative. Furthermore, the one experiment showing mitochondrial complex I inhibition utilized a 100 μM dose of CPC, which, although exposure was brief (<10 min), is a concentration several-fold higher than the amount known to cause significant cytotoxicity to mammalian cells in 1 hr (Raut *et al*., 2022). This finding raises the question of the levels of cell death accompanying the reported mitotoxicity (Datta *et al*., 2017b).

A third study used primary cells and very high doses of CPC: 100 μM CPC caused a brief OCR stimulation (in the first 5-10 min), followed by a 40% reduction in OCR in toad bladder cells after 60 minutes exposure (Saladino *et al*., 1971). These OCR experiments were conducted in the presence of 5 mM glucose, conditions under which ample glycolysis can operate to produce ATP independent of mitochondria, which may complicate data interpretation. By the utility of light and electron microscopy combined with time-lapse cinemicrography, they also found mitochondrial deformation after just 5 min of exposure at the same CPC concentration, while they also observed cell swelling by 30 minutes, suggesting accompanying cytotoxicity (Saladino *et al*., 1971). Electron microscopy is a powerful method to reveal mitochondrial ultrastructure but requires heavy sample processing which may convolve with toxicant effects on structure to complicate interpretation of toxicant data. Altogether, the limited pool of eukaryotic studies and mitochondrial toxicity studies necessitates further research into the toxicity of CPC.

These important studies (Saladino *et al*., 1971; Chávez and Concepcion, 1982; Datta *et al*., 2017b) suggest that CPC is a mitochondrial toxicant, yet outstanding questions remain, such as whether CPC affects mitochondria in living human and other mammalian cells including primary cells and cells that are not inherently susceptible to mitotoxicity. What is needed are exposures at non-cytotoxic and exposure-relevant CPC doses and under conditions (e.g., galactose media) designed to highlight mitochondrial effects and their mechanism(s). In the current study, we have compared CPC’s mitotoxicity to its immunotoxicity and have investigated CPC’s mitochondrial mechanisms of action and its nanostructural effects using super-resolution microscopy.

As part of a larger study of CPC inhibition of immune cell signaling (Raut *et al*., 2022), we unexpectedly discovered CPC mitotoxicity, the topic of this manuscript. We previously discovered that CPC inhibits function (degranulation) of the immune cell type mast cells (Raut *et al*., 2022), a process which is ATP-dependent (Burgoyne and Morgan, 2003). Mast cells are enriched at environmental interfaces (Kuby, 1997; Theoharides *et al*., 2012; Blank and Benhamou, 2013), so they are poised for exposure to CPC via inhalation, food ingestion, and product application. Found in most tissues (Dvorak, 1986; Theoharides and Sant, 1991; Blank and al., 2007; Farrell and al., 1995) even the brain (Silver and Curley, 2013), mast cells play key roles in both nervous-system (Theoharides *et al*., 2012; Theoharides *et al*., 2015; Theoharides *et al*., 2016; Theoharides *et al*., 2019) and immune-related functions (Metcalfe *et al*., 1997; Galli *et al*., 2005; Dawicki and Marshall, 2007; Sinnamon *et al*., 2008; Ribatti and Crivellato, 2009; Khazaie *et al*., 2011; Pittoni *et al*., 2011; Ribatti and Crivellato, 2011; Anand *et al*., 2012;

Krystel-Whittemore *et al*., 2015; Johnzon *et al*., 2016). The cell line utilized for this study to model human mast cells was the rat basophilic leukemia (RBL-2H3) cell line. RBL-2H3 cells express the high-affinity immunoglobin E (IgE) receptor, FcεRI and serve as an appropriate human mast cell model (Falcone *et al*., 2018) as they share similarities to signaling pathways found in humans (Mohr and Fewtrell, 1987). Other advantages of RBL-2H3 cells for toxicology studies are their rapid doubling rate (Gray *et al*., 2015) and the similarity of their biochemical responses to those of the canonical mast cell model primary bone marrow-derived mouse mast cells (Zaitsu *et al*., 2007; Thrasher *et al*., 2013; Alsaleh *et al*., 2016). Mast cell function within the body hinges on their ability to undergo degranulation, which is an antigen (Ag)-dependent process that triggers the release of granules containing bioactive chemicals.

It is important to assess the CPC exposure in multiple cell types considering the many opportunities for location of CPC exposure because of the variety of CPC-containing products. Primary human keratinocytes are a predominant cell type in epidermal tissue and thus are a model for cells at interfaces between the human body and the environment through which exposure to CPC occurs. Another cell type used to confirm effects of CPC exposure is the immortalized NIH-3T3 cell line, mouse embryonic fibroblasts. All three cell types (NIH-3T3, primary human keratinocytes, and RBL-2H3) used in this study were previously utilized to evaluate mitotoxicity of antibacterial agent triclosan (TCS) (Weatherly *et al*., 2016; Weatherly *et al*., 2018).

CPC cytotoxicity has previously been assessed in NIH-3T3 and RBL-2H3 cells. CPC (≤ 15 μM) was found to be non-cytotoxic in NIH-3T3 and RBL-2H3 cells via trypan blue-exclusion assay and confirmed via lactate dehydrogenase assay for RBL-2H3 cells (Raut *et al*., 2022). In the current study, we have conducted further cytotoxicity tests to ensure that all dosing regimens used do not cause cell death. All experiments conducted in this study used doses ≤ 10 μM CPC, which were non-cytotoxic to these three cell types during the timing and buffer conditions utilized.

In this study, super-resolution imaging of live cells (Hess *et al*., 2007) via FPALM (Hess *et al*., 2006) was employed to visualize the nanoscale details of mitochondrial structure, with the Dendra2TOM20 outer mitochondrial membrane marker. FPALM breaks the diffraction limit of conventional microscopy resolution by a factor of ∼10X, helpful in mitochondrial studies given that key mitochondrial features are smaller than the ∼250 nm diffraction limit that causes blurring and obscuring of fine details. The advantage of FPALM over traditional electron microscopy is the ability to image living cells and to do so without the heavy chemical and physical processing required by electron microscopy, which may confound interpretation of toxicant effects. FPALM works by employing photoactivatable fluorophores within a sample and activating a subset of those fluorophores, localizing and recording their central positions, then photobleaching them and repeating the cycle. When data are integrated over numerous molecules, a high-resolution image of the sample emerges (Hess *et al*., 2006). To the best of our knowledge, our previous usage of FPALM to elucidate 2-and 3-D nanoscale mitochondrial structural toxicity of antibacterial agent triclosan are among the first usages of super-resolution microscopy in the field of toxicology (Weatherly *et al*., 2018, Sangroula *et al*., 2020). Moreover, a current search of years of Society of Toxicology annual meeting abstracts yields no uses of FPALM or other super-resolution microscopy methods, other than our own–highlighting the novel use of this method in the field of toxicology.

Effects of non-cytotoxic levels of CPC on mitochondria in intact mammalian and primary human cells, as well as mechanistic aspects, are not yet known. Thus, we aim to discover CPC effects specifically on mitochondrial Ca^2+^ via our development of a new plate reader-based mitochondrial-Ca^2+^ assay and nanostructural changes by the novel use of FPALM. Here, we examined the effects of CPC in human, rat, and mouse cells on mitochondrial function: ATP production, oxygen consumption, mitochondrial Ca^2+^ buffering, and nanostructural integrity. Our work demonstrates that CPC impairs mitochondrial function at doses as low as 3000-fold lower than the dosages currently used in personal care and food products.

## Methods

### Chemicals and Reagents

Cetylpyridinium chloride (CPC; 99% purity, VWR; CAS no. 123-03-5) was prepared, including concentration determination, as previously detailed in aqueous solution (Raut *et al*., 2022), using a method that avoids organic solvents that could cause adverse cell effects. For the mitochondrial-Ca^2+^ assay, CPC was dissolved into Tyrode’s buffer (made with either galactose or glucose) (Hutchinson *et al*., 2011), then bovine serum albumin (BSA; 1 g/L) was added as in (Raut *et al*., 2022); this is called BT. For all other experiments, CPC was dissolved into cell culture water (CCW; VWR) as a base for preparation of ToxGlo Media, which was used in all experiments other than mitochondrial Ca^2+^. CPC does not absorb UV–Vis light beyond approximately 280 nm (Bernauer *et al*., 2015) and thus will not interfere via absorption with probes used in this study (which all use excitation wavelengths of 360 nm or higher, as noted in sections below).

### Preparation of ToxGlo Media with Glucose or Galactose

CPC solution prepared in CCW was combined with additional ingredients including L-glutamine, to create ToxGlo Media with 5.6 mM glucose or galactose, using the published recipe (Weatherly *et al*., 2016). Glucose or galactose ToxGlo Media was made fresh on each day of experimentation with the final component BSA added, then media were brought to pH 7.4. The pH of 0 μM CPC control media was measured first, to avoid CPC cross-contamination.

### Cell Culture

#### RBL-2H3 Cell Culture

RBL-2H3 mast cells were cultured as described (Hutchinson *et al*., 2011).

#### NIH-3T3 Cell Culture

NIH-3T3 mouse fibroblast cells (ATCC) were cultured as described (Curthoys *et al*., 2019).

#### Primary Keratinocyte Culture

Primary adult human epidermal keratinocytes (Lifeline Cell Technology) were cultured in DermaLife media as per manufacturer’s instructions.

### Cytotoxicity and ATP Production Assay

Cytotoxicity and ATP Production were measured with the use of a Mitochondrial ToxGlo™ Assay Kit (Promega), as described in (Weatherly *et al*., 2016), adapted for CPC. RBL-2H3 cells, NIH-3T3 cells, or primary human keratinocytes were assayed. For the 60 min CPC treatments ahead of cytotoxicity assessment, pre-warmed CPC at 2X concentration in ToxGlo Media was added to cells, resulting in 1X [CPC] dissolved in glucose or galactose ToxGlo Media for cell exposures. Samples serving as positive controls for the cytotoxicity measurement received 72 μg/mL membrane-permeabilizer digitonin (Promega) in respective ToxGlo Media type (glucose or galactose). Following this 60 min treatment at 37℃/5% CO_2_, cytotoxicity was measured as in (Weatherly *et al*., 2016). Validity of the assay was determined by digitonin wells’ fluorescence values ≥ 2x of those of control wells. Cells were next incubated with ATP reagent at room temperature for 5 min before luminescence was measured. All replicates were normalized to 100% of the average of the untreated control (0 μM CPC) on a given experimental day.

### Oxygen Consumption Rate Assay

Oxygen consumption rates of RBL-2H3 mast cells and of primary human keratinocytes were assessed via the Oxygen Consumption Rate Assay Kit (Cayman Chemical Company) as published (Weatherly *et al*., 2016). The same glucose-free, galactose ToxGlo Media as in the Mitochondrial ToxGlo Assay was made on the second day of experimentation and used to prepare CPC treatments. Quadruplet cell samples were assayed for each of the following four conditions: CPC with MitoXpress-Xtra probe, CPC without probe, no CPC with probe, and no CPC without probe. Probe-containing wells received 2.38% (v/v) of the probe. Foil plate covers (Zymo Research) were used to seal wells from the ambient environment and to eliminate oxygen entry into the wells. Phosphorescence measurements were taken for 3 hr at 10 min intervals at 37°C, 360 ± 40 nm excitation, 645 ± 15 nm emission, gain 90, bottom reading, and at normal speed.

The average of phosphorescence values from wells containing no probe was subtracted from the average values of corresponding wells containing probe; this process was repeated for each CPC dose and time point. The 30-180 min data points were utilized, to avoid initial (t = 0, 10, 20 min) phosphorescence drift due to plate warming. A linear regression line was fit to the 30-180 min data and was plotted, in order to determine oxygen consumption rate (raw phosphorescence units per minute; RFU/min). Accepted experiments met the criteria of having a control (“0 μM CPC”) with 1.) an RFU increase > 60 and 2.) an R^2^ linear regression fit value > 0.75. Areas under the curve were calculated with GraphPad Prism for each experiment, normalized to control (“0 μM CPC”), and then averaged across experiments.

### RBL-2H3 Cell Degranulation Assay

Degranulation assay was performed as detailed in (Weatherly *et al*., 2013) adapted for use with CPC with protocol modifications as follow: CPC treatments (60 min) were delivered in ToxGlo Media with glucose or galactose, and DNP-BSA antigen was administered at 0.001 μg/mL.

### Mitochondrial Ca^2+^ assay with Genetically Encoded Indicator CEPIA2mt

RBL-2H3 cells were transfected with pCMV CEPIA2mt construct, a gift from Masamitsu Iino (Addgene plasmid # 58218; http://n2t.net/addgene:58218 ; PRID:Addgene_58218) (Suzuki *et al*., 2014) via an RBL-2H3-specific Amaxa Nucleofector Transfection Kit T (Lonza), as done in (Weatherly *et al*., 2018). The construct was utilized to measure levels of mitochondrial Ca^2+^ in response to CPC treatment. “Mock” transfected cells underwent the same electroporation process but without plasmid DNA. Following electroporation, cells were added to 200 μL phenol red- free RBL media at 100,000 cells/well in a black, clear-bottom, tissue culture-treated, 96- microwell Greiner Bio-One® plate and grown at 37℃/5% CO_2_ overnight. The next day, cell media was dumped, and cells were washed twice with fresh control (0 μM CPC) BT. The control BT used was made as noted in “Chemicals and Reagents,” with the glucose (5.6 mM) substituted for galactose (5.6 mM or 10 mM) in a series of experiments to assay the difference in mitochondrial Ca^2+^ based on cellular ability to perform glycolysis. The Mock-transfected cell wells, the “No CPC” cell wells, and “60 min CPC” exposed cells then received 200 μL control BT while “90 min CPC” exposed cells received 200 μL (5 or 10 μM) CPC in BT. The plate was incubated in the 37℃/5% CO_2_ incubator for 30 min. Following this incubation, the plate was dumped, and cells received 200 μL (5 or 10 μM) CPC (“90 min CPC” and “60 min CPC” exposed cells) or control BT without CPC (Mock and “No CPC”). Cell fluorescence was immediately read for 60 min at 485 ± 20 nm excitation, 528 ± 20 nm emission, gain 120, bottom reading, and at normal speed for 42 sec intervals. The average fluorescence of “Mock” transfected cells (which is average background fluorescence from the cells, the plate, and the BT) was subtracted from each curve at each respective time point from those of different treatment groups. Additionally, transfection using plasmid preparations of differing concentrations of DNA with differing levels of DNA purity adds variation to the total amount of transfection and subsequent fluorescence of the CEPIA2mt construct. Accepted experiments met the criterion that the “No CPC” group had 0 min, “Mock”-subtracted raw fluorescence values (RFU) of > 4,000; this criterion indicated that reporter construct transfection levels were sufficient. Areas under the curve were calculated with GraphPad Prism for each experiment, normalized to control (“No CPC”), and then averaged across experiments.

### Fluorescence Photoactivation Localization Microscopy (FPALM) Imaging and Processing

NIH-3T3 cells were grown in complete NIH-3T3 media (DMEM with 4.5g/L Glucose and 4mM L-Glutamine (Lonza), 10%, iron-fortified Bovine Calf Serum (Sigma Aldrich), Penicillin (100 I.U./mL)-Streptomycin (100 μg/mL) (ATCC). Solutions were made in advance using Millipore Stericups. Cells were plated into 35 mm MatTek dishes (35 mm, coverglass diameter 20 mm) with complete NIH-3T3 media (without phenol red and without any antibiotics) and grown in 80,000 cells/MatTek dish for ∼24h in 37℃/5% CO_2_ cell culture incubator. Then confluent NIH-3T3 cells were transfected with 1 μg Dendra2Tom20/MatTek dish in OptiMEM (reduced serum medium, with L-glutamine, without phenol red, Gibco) and Lipofectamine 3000 and incubated for 6 hours at 37℃/5% CO_2_. After incubation, the previous OptiMEM media/DNA mixture was removed and replaced with new complete NIH-3T3 media (without phenol red and without any antibiotics) and incubated for 12-16 hours at 37℃/5% CO_2_ until ready for imaging. Prior to imaging, media was removed, and cells were washed with pre-warmed control ToxGlo galactose media and pre-exposed with either 0 µM CPC or 5 µM CPC in ToxGlo galactose media for 60 minutes, at 37LJ/5% CO_2_.

After 60 minutes of exposure to either 0 µM CPC or 5 µM CPC media, imaging was performed such that 5 cells were imaged per MatTek dish for up to 30 minutes after removal from the incubator. Cells were imaged on an Olympus IX71 microscope via widefield laser illumination (Hess *et al*., 2006) to allow imaging of mitochondria within the cytoplasm of the cell. A 558 nm laser (CrystalLaser LC, CL558-100) was utilized to both photoswitch and photoactivate the Dendra2 fluorophore attached to TOM20; this provided a better signal to noise ratio than when using a separate activation beam. Methods used for single-color super-resolution imaging and localization follow previously published methods (Hess *et al*., 2006; Weatherly *et al*., 2018; Curthoys *et al*., 2019) using temporal median background subtraction (Piccardi, 2004).

The experimental setup was similar to that used previously (Weatherly *et al*., 2018). Here, we provide a detailed list of components since there are some minor differences.

### Illumination Path for Cell Selection

Cells were chosen based on observation of intracellular green fluorescence under excitation of inactive Dendra2 by a mercury lamp light passed through a beam expander (BE10 10X, ThorLabs), an excitation bandpass filter (476/10, Noran Microscopes), and a +300 mm lens. After collection by the 60X 1.45NA oil objective, the Dendra2 fluorescence was bandpass-filtered (ET525/50 m, Chroma) before reaching the oculars. After switching off the lamp, changing to the laser acquisition filter set, and opening the laser shutter, the fluorescence from the photoactivated Dendra2 molecules was brought into focus within a few seconds and then acquisition was initiated.

### Illumination Pathway for FPALM Acquisition

A 558 nm laser focused with a convex lens (f = +350 mm, Newport Corporation) into the back aperture of an inverted microscope (1X71, Olympus) and through a 60X 1.45 NA oil objective lens (Olympus) to yield 23.7mW total power and an average intensity of 6.3 kW/cm^2^ at the sample. Fluorescence emission from Dendra2TOM20 molecules was captured through the same objective, passed through a dichroic (405/488/561/635 nm multiband, Semrock), two notch filters at a wavelength of 561 nm (Semrock), and one notch filter at a center wavelength of 405 nm (Semrock) to reduce laser transmission. A mechanical aperture at the focal point of the tube lens was used to block light outside the region of interest, and the remaining light passed through a telescope composed of two achromatic lenses (f = +200 mm and f = +400 mm, Newport Corporation) for additional magnification which resulted in a measured camera pixel size equivalent to about 109 nm at the sample. A bandpass filter (HQ590/75 M, Chroma) filtered out potential background fluorescence, and the resultant image was detected by the sensor of an EMCCD (iXon+ DU897DCS-BV, Andor Scientific) with an EM gain of 200 and an exposure time of 10.79 ms per frame. A set of 10,000 frames was collected from each region, imaged, and stored as a “.tif” stack. The use of the 558 nm beam as both the readout and activation laser provided a better signal to noise ratio in this setup than including the use of a 405 nm activation beam, and as sufficient localizations were detected, this was the only beam used.

### FPALM Data Analysis

Data collected by the EMCCD camera were analyzed using scripts written for single color localization and rendering (Hess *et al*., 2006; Gudheti *et al*., 2013; Curthoys *et al*., 2019) using MATLAB (MathWorks Inc.). Another script was used to select a subset of the generated renders, cropping to keep a central, rectangular area and to exclude regions beyond the imaging region. Using ImageJ software (NIH), an algorithm was run to repetitively apply a smoothing function for each image to create a continuous shape from individual localizations for analysis of size and shape parameters corresponding to the structures which the localizations represented. ImageJ was then utilized to generate an intensity-based constant-threshold mask to include data from the smoothed images, with the same threshold for all images. After converting pixels to appropriate units for the parameters, data were imported into GraphPad Prism for graphing and statistical analysis.

### Donut Counting Analysis

Unmodified renders of the localization data of both the 0 µM CPC and 5 µM CPC samples were combined into a folder and a MATLAB script was used to randomize the file names. The set of randomized images were sent out to the two researchers who collected the data, as well as a third, who had not seen the images before. The three were asked to count the number of donut-like spherical shapes observed in each of the images. The files were then mapped back to the original images with unmodified file names to determine the counts for each of the 0 µM CPC and 5 µM CPC samples. The counts were averaged for each of the images, which typically consisted of 1 cell, and were plotted into GraphPad Prism.

### Fourier Transform Analysis

A MATLAB script was written to process the unmodified renders of localization data and quantify the morphological differences observed in the donut analysis. Image files were read, converted into a Fast Fourier Transform (FFT), and averaged separately for both the 0 µM CPC and 5 µM CPC groups. The FFT amplitude was then thresholded with both a lower and upper bound to exclude spatial frequencies which were beyond the FPALM resolution limit or otherwise outside of the length scales corresponding to the relevant image features. These threshold values are estimates, as the prevalence of features at different length scales is not consistent between the samples, but it is this observed variance between 0 µM CPC and 5 µM CPC samples that is of interest.

The thresholded FFT images were analyzed to obtain the distance from the origin (zero spatial frequency) to each point which was between the two amplitude thresholds. A rectangular region spanning the x-axis (±5 pixels above or below the origin) was taken, and similarly, this was done along the y axis (±5 pixels to the left or right of the origin). The cumulative histogram of the data contained as a function of distance along the x and y axes, respectively, was then determined. The ratio of these two histograms was plotted against the distance from the origin; a ratio of approximately one suggested the data was not directionally biased, while a value larger or smaller than one suggested features more prevalent in the x or y direction, respectively, at the given distance. The ratios asymptotically reached a value of one at large distances, since all of the FFT features were contained by the region when extended to a very large distance from the center.

### Statistical Analyses

Using GraphPad Prism, statistical significance was determined by one-way ANOVA with a Tukey’s post-hoc test for the assays of cytotoxicity/ATP production, oxygen consumption rate (also one-tailed paired t-test), degranulation, mitochondrial Ca^2+^, and changes in mitochondrial shape extracted from FPALM images. ATP inhibition EC_50_ was also determined by Prism software. The Mann-Whitney test was used for donut counting data in GraphPad Prism. The plots generated from the FFT data were compared using a Kolmogorov-Smirnov test.

## Results

### CPC potently inhibits ATP production in RBL-2H3, NIH-3T3, and primary human keratinocyte cells

To determine the effect of CPC on cytotoxicity and mitochondrial ATP production, we utilized the Mitochondrial ToxGlo Assay kit (Promega). Cells were plated on the day of experimentation and received varying concentrations of CPC. A digitonin-lysed cell triplicate was added as a positive control to validate the cytotoxicity assay. Following CPC incubation, the bis-AAF-R110 substrate for fluorescent measurement of cytotoxicity was added. CPC concentrations of 10 μM and lower are known to be non-cytotoxic to RBL-2H3 and NIH-3T3 cells for 60 minutes via lactate dehydrogenase and trypan blue exclusion assays (Raut *et al*., 2022), but, due to the 90 minute total incubation time and differences in cell media used for this assay, this cytotoxicity assay component was necessary to support the ATP results. Following the fluorescent reading, a cell-lysing ATP reagent was added to produce a luminescent signal that is proportional to the amount of ATP present.

In galactose media, the EC_50_ of ATP production when RBL-2H3 cells were exposed to CPC for 90 min was 1.7 μM, via GraphPad Prism statistical analysis (Figure 1A). Although at 10 μM CPC, levels of ATP in glucose media were 88% ± 3% (SEM) of the control (Figure 1A), no statistically significant decrease in ATP was found due to CPC exposure in the presence of glucose. This difference between CPC effects on ATP production in galactose vs. glucose media indicates that CPC’s effect is on mitochondrial ATP production, specifically–indicating that CPC is a mitochondrial toxicant. In these RBL-2H3 cell experiments, no statistically-significant effect on cytotoxicity was seen across media type (glucose or galactose) or concentration of CPC assayed for 90 min.

**Figure 1.**
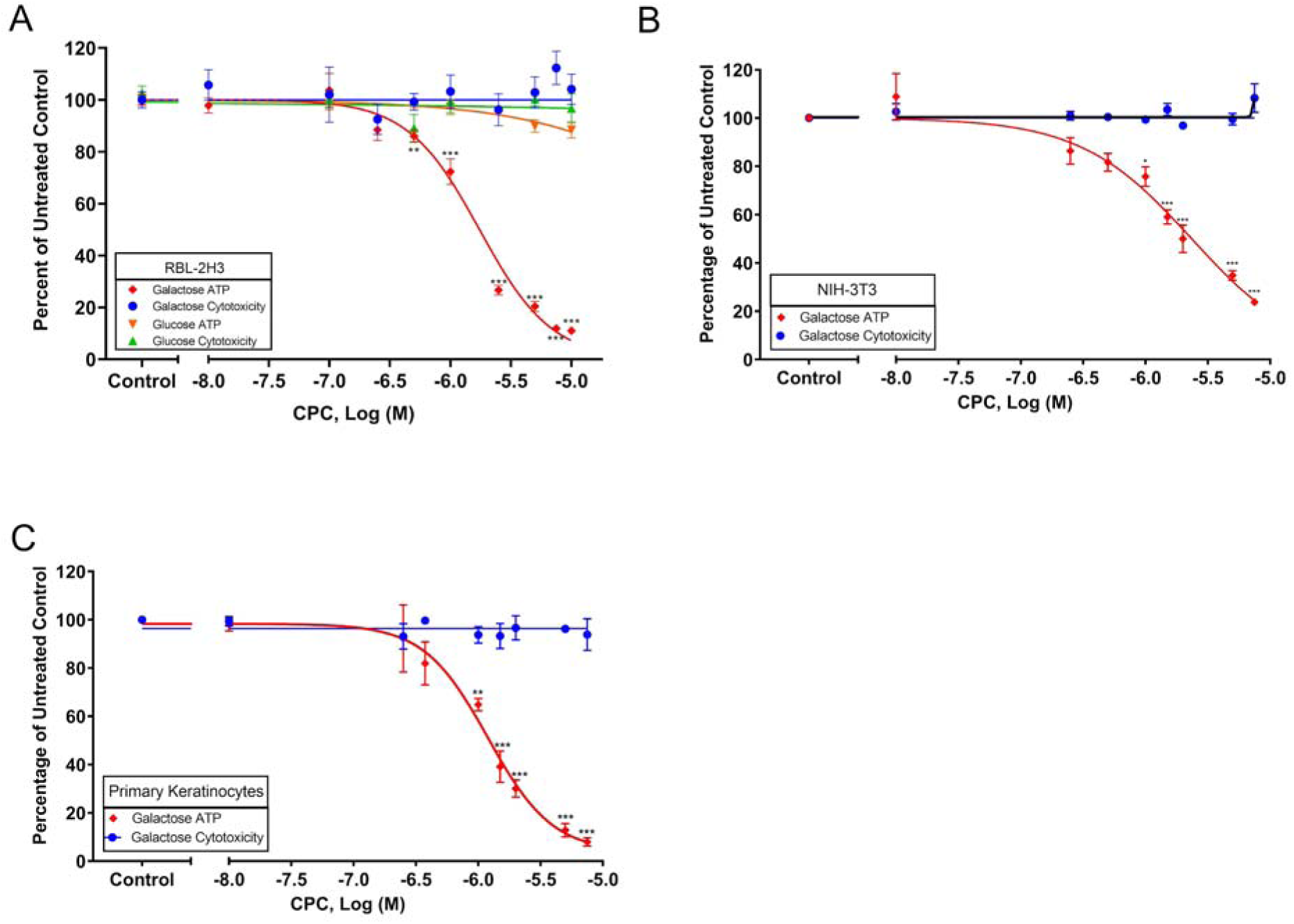
Effects of CPC on RBL-2H3 mast cell, NIH-3T3 mouse fibroblast, and primary human keratinocyte cytotoxicity and ATP production in glucose and galactose media. RBL-2H3 mast cell assay was carried out in glucose or in glucose-free, galactose-containing media **(A)**, whereas NIH-3T3 mouse fibroblast **(B)** and primary human keratinocyte **(C)** assays were conducted solely in galactose media, to cause cells to produce ATP in mitochondria. All cell types were exposed to varied CPC doses for 90 min prior to cytotoxicity measurements. Fluorescence and luminescence values were normalized as percentages of untreated control average values and shown as mean ± SEM. Figures are derived from at least three independent experiments with duplicates or triplicates for each CPC dosage. Statistically significant results indicated by *p < 0.05, **p < 0.01, ***p < 0.001 compared to control (0 μM CPC), one-way ANOVA followed by Tukey’s post-hoc test.

We confirmed that CPC had no significant effect on the fluorescence or luminescence signals of the Mitochondrial ToxGlo assay, confirming that the observed CPC effects on ATP production are due to a true cellular effect (Supplemental Figures S1 & S2).

To test whether this CPC mitotoxicity is a universal effect rather than one specific solely to rat mast RBL-2H3 cells, ATP production was also measured in NIH-3T3 mouse fibroblasts and in primary human keratinocytes. While in galactose media, the EC_50_ of ATP production of NIH-3T3 cells exposed to CPC for 90 min was 10 μM (Figure 1B) and in primary human keratinocytes was 1.2 μM (Figure 1C). CPC did not cause cytotoxicity under the tested conditions in both NIH-3T3 (Figure 1B) and primary human keratinocytes (Figure 1C). At 10 μM CPC, levels of ATP in galactose media decreased 76% ± 1% (SEM) in NIH-3T3 cells (Figure 1B), 92% ± 2% (SEM) in primary human keratinocytes (Figure 1C), when compared to the control. Overall, these data suggest that CPC is a mitochondrial toxicant in various cell types, from three species, including a primary cell type.

### CPC dampens O_2_ consumption rate in primary human keratinocytes and in RBL-2H3 cells

To determine the effect that CPC has on O_2_ consumption rate (OCR), the OCR kit from Cayman Chemical Company was used. This kit relies on the use of a phosphorescent probe that is quenched by O_2_; thus, O_2_-consuming mitochondrial respiration reduces O_2_ over time, leading to an increase in phosphorescence reading. We previously developed and validated this assay with RBL-2H3 cells (Weatherly *et al*., 2016). To validate the experiments, we assayed CPC effects on the experimental signal (Supplemental Figures S3-S5). CPC does not change the background (no-probe) phosphorescence of samples with cells (Supplemental Figure S3). CPC does not change the OCR probe’s phosphorescence of samples without cells (Supplemental Figure S4), showing that the OCR cellular data (Figure 2) is truly due to CPC effects on cellular oxygen consumption rates. Additionally, we confirmed that CPC does not interfere with the phosphorescent probe signal at the start of the OCR experiments, with or without cells (Supplemental Figure S5).

**Figure 2.**
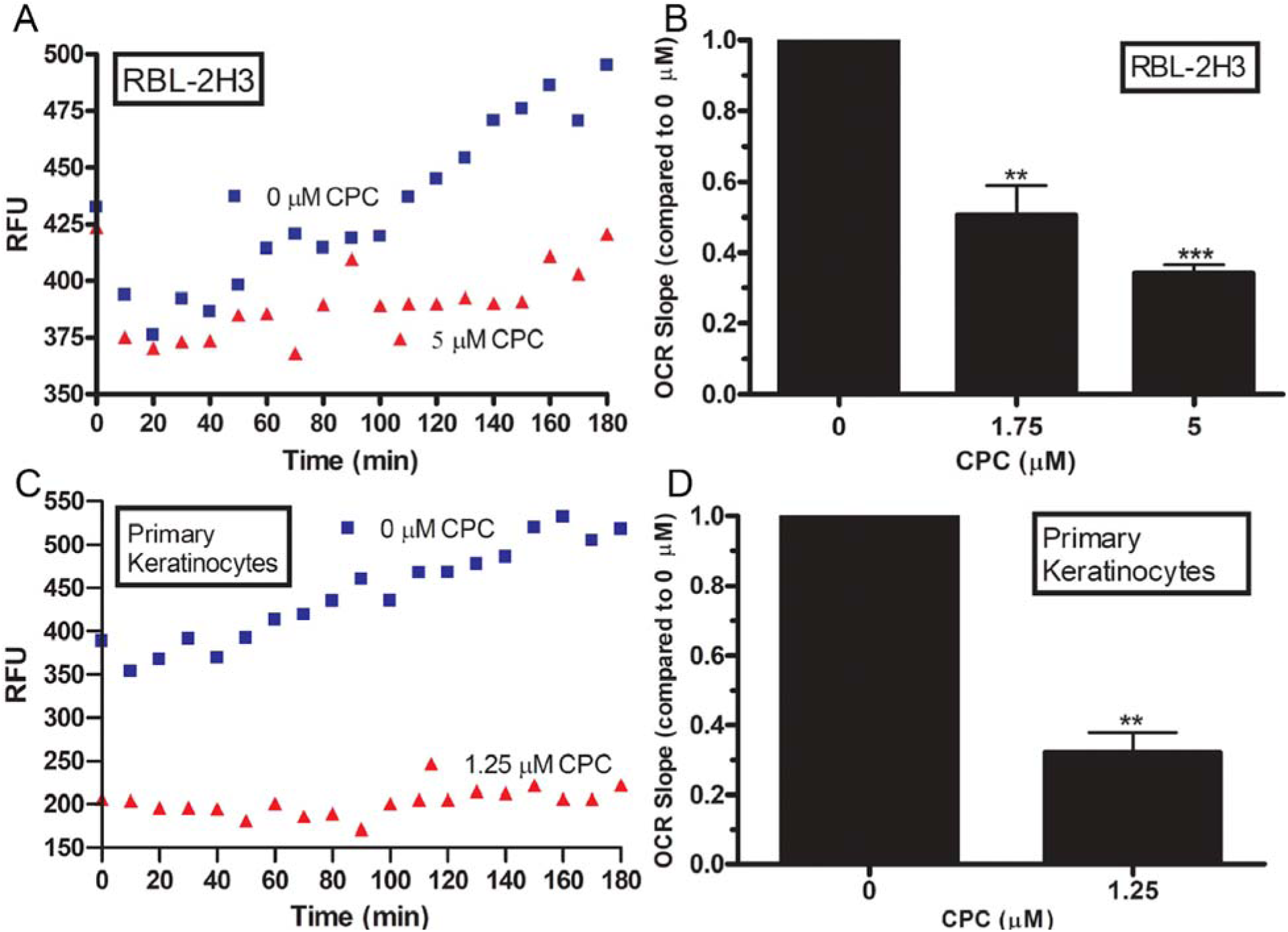
Oxygen consumption rate in CPC-treated RBL-2H3 cells and primary human keratinocytes in galactose media. OCR was measured in cells exposed to CPC for 3 hr with data fit to a linear line, with best-fit slope (RFU/min) based on 10 min incremental measurements (t = 30 to t = 180). Representative graph of RBL-2H3 OCR data **(A)**. Slope linear line of best fit for 1.75 μM and 5 μM CPC normalized to untreated (0 μM CPC) control, shown as mean ± SEM, **p < 0.01 and ***p < 0.001, determined by one-way ANOVA by Tukey’s post-hoc test. **(B)** Representative graph of primary human keratinocyte OCR data **(C)** Slope of linear line of best fit for 1.25 μM normalized to untreated (0 μM CPC) control, shown as mean ± SEM, **p < 0.01, determined by one-tailed paired t-test with 95% Confidence Interval **(D)**. Data gathered from three experiments, each with at least three replicates per sample.

The averaged, background-subtracted value for RBL-2H3 cells was plotted over time and shown as a representative figure (Figure 2A). A reduced rate of increase in phosphorescence over time is seen with CPC, when compared to the 0 μM control line (Figure 2A). The slopes of the lines which represent the OCR, in the 0 μM, 1.75 μM, and 5 μM CPC groups from t = 30 min to 180 min, were determined and normalized to that of the 0 μM control group (Figure 2B). The OCR for the 5 μM CPC-treated RBL-2H3 cells decreased to 34% ± 2% (SEM) of the untreated control, representing roughly a two-thirds decrease in oxygen consumption, while that of the 1.75 μM CPC-treated was 51% ± 8% (SEM), representing roughly a one-half decrease in oxygen consumption (Figure 2B). The averaged, background-subtracted value for primary human keratinocyte was plotted over time and shown as a representative figure (Figure 2C). The slope of the line in the 0 μM and 1.25 μM CPC groups from t = 30 min to 180 min were determined and normalized to that of the 0 μM control group (Figure 2D). The OCR for the 1.25 μM CPC-treated primary human keratinocytes was 32.1 % ± 0.1% (SEM) of the untreated control, representing a roughly two-thirds decrease in oxygen consumption (Figure 2D).

### CPC inhibits RBL-2H3 degranulation under mitochondrial ToxGlo assay conditions

To assess differences in CPC’s ability to inhibit RBL-2H3 cells degranulation under ToxGlo assay conditions, the RBL-2H3 degranulation assay (Figure 1 in Raut, Weller *et al*., 2022) was repeated without a pre-incubation. The dose of antigen (Ag) utilized, 0.001 μg/mL, was chosen because it elicited a moderate absolute degranulation response when compared to the maximal possible granule release when in the absence of CPC: absolute degranulation in glucose media was 32% ± 4% (SEM), and in galactose media was 23.4% ± 0.5% (SEM).

There were no significant changes in levels of CPC suppression of degranulation between the glucose and galactose groups: levels of degranulation at 10 μM CPC were reduced to 51% ± 3% (SEM) of the control value when in glucose media and to 57% ± 2% (SEM) of the control value in galactose media (Figure 3). In both media types, inhibition of degranulation was dose-responsive, with significance beginning at 1 μM for the cells in glucose media and at 5 μM for those in galactose media (Figure 3). Previously, we found that 10 μM CPC inhibits degranulation to 30% ± 6% (SEM) of control values when in glucose-BT buffer (Raut *et al*., 2022), indicating that CPC causes less inhibition of degranulation under the ToxGlo assay conditions. In a previous study (Raut *et al*., 2022), we showed that CPC inhibition of degranulation is a true cellular effect, not an interference of CPC with the enzymatic reaction and fluorophore used to assess degranulation.

**Figure 3.**
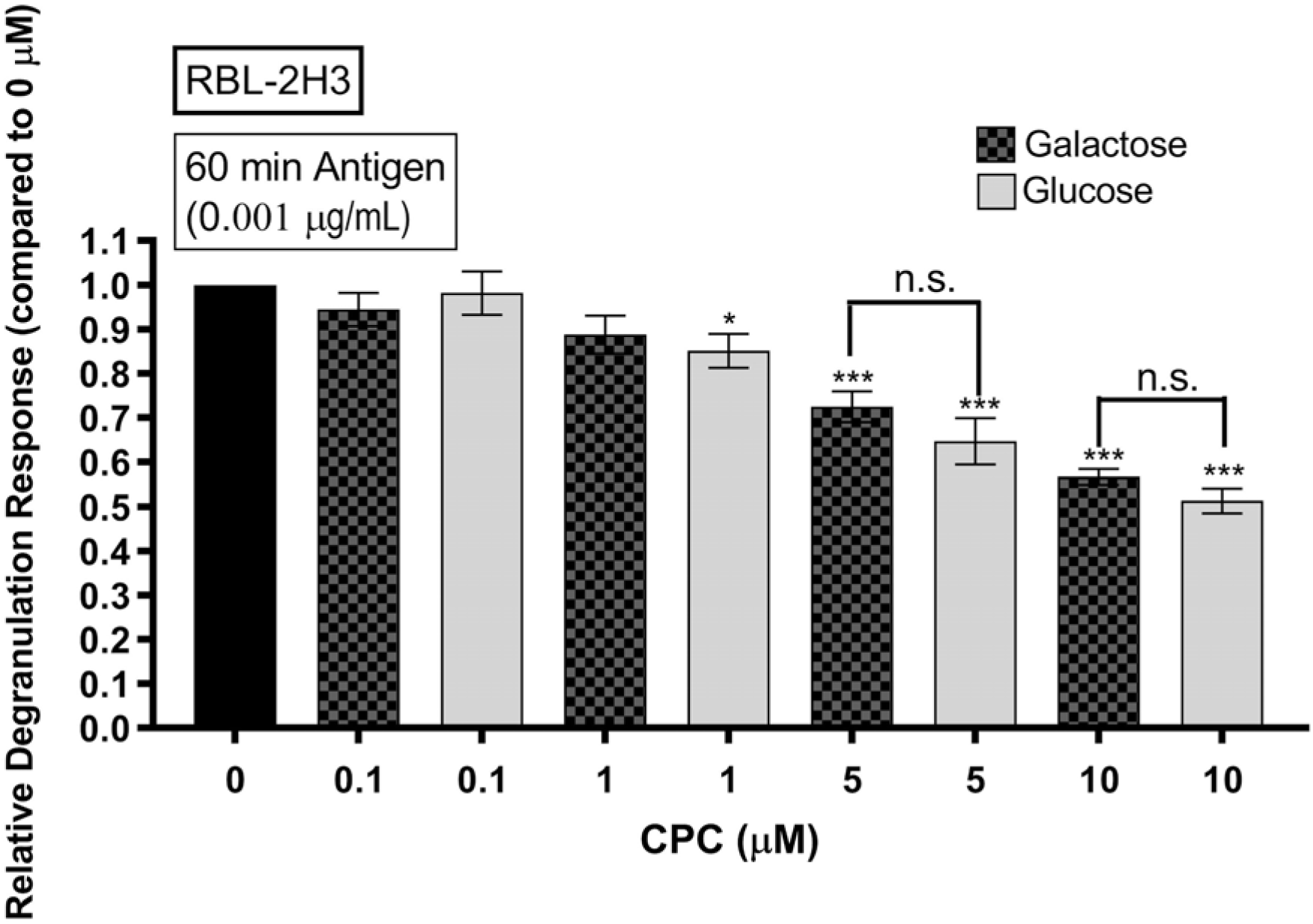
Relative degranulation levels of RBL-2H3 cells after CPC treatment in glucose and galactose media. Cells primed with IgE for 60 min followed by treatment with CPC + 0.001 μg/mL antigen (Ag) for 60 min. CPC concentrations ranged between 0 and 10 μM. Degranulation values were normalized to control (0 μM CPC + Ag). Data shown as mean ± SEM of three independent experiments with triplicates of each sample in each experiment. n.s. = not significant, *p < 0.05 and ***p < 0.001, determined by one-way ANOVA followed by Tukey’s post-hoc test.

### CPC lowers mitochondrial Ca^2+^ levels in galactose-containing buffer

To assess whether CPC affects mitochondrial Ca^2+^ levels, RBL-2H3 cells were transfected with the mitochondrial-targeted, fluorescent construct CEPIA2mt and tested in a novel plate reader-based assay that allows for relatively rapid assessment of numerous treatment conditions. CPC addition led to decreased reporter fluorescence levels, under a variety of exposure conditions: both 5 μM (Figure 4A) and 10 μM (Figure 4B-4C) CPC concentrations; both “60 min CPC’’ and “90 min CPC” exposure timings (Figures 4A-4C); and in both 5.6 mM (Figures 4A-4B) or 10 mM (Figure 4C) galactose buffer. In “90 min CPC” representative graphs (Figures 4A-4C), refilling of Ca^2+^ into the mitochondrial matrix is apparent at the later time points. AUC analyses of the galactose-buffer groups revealed statistically significant decreases in reporter fluorescence due to CPC under all conditions tested (various CPC timing and dosing and galactose concentrations as indicated in Figure 4D).

**Figure 4.**
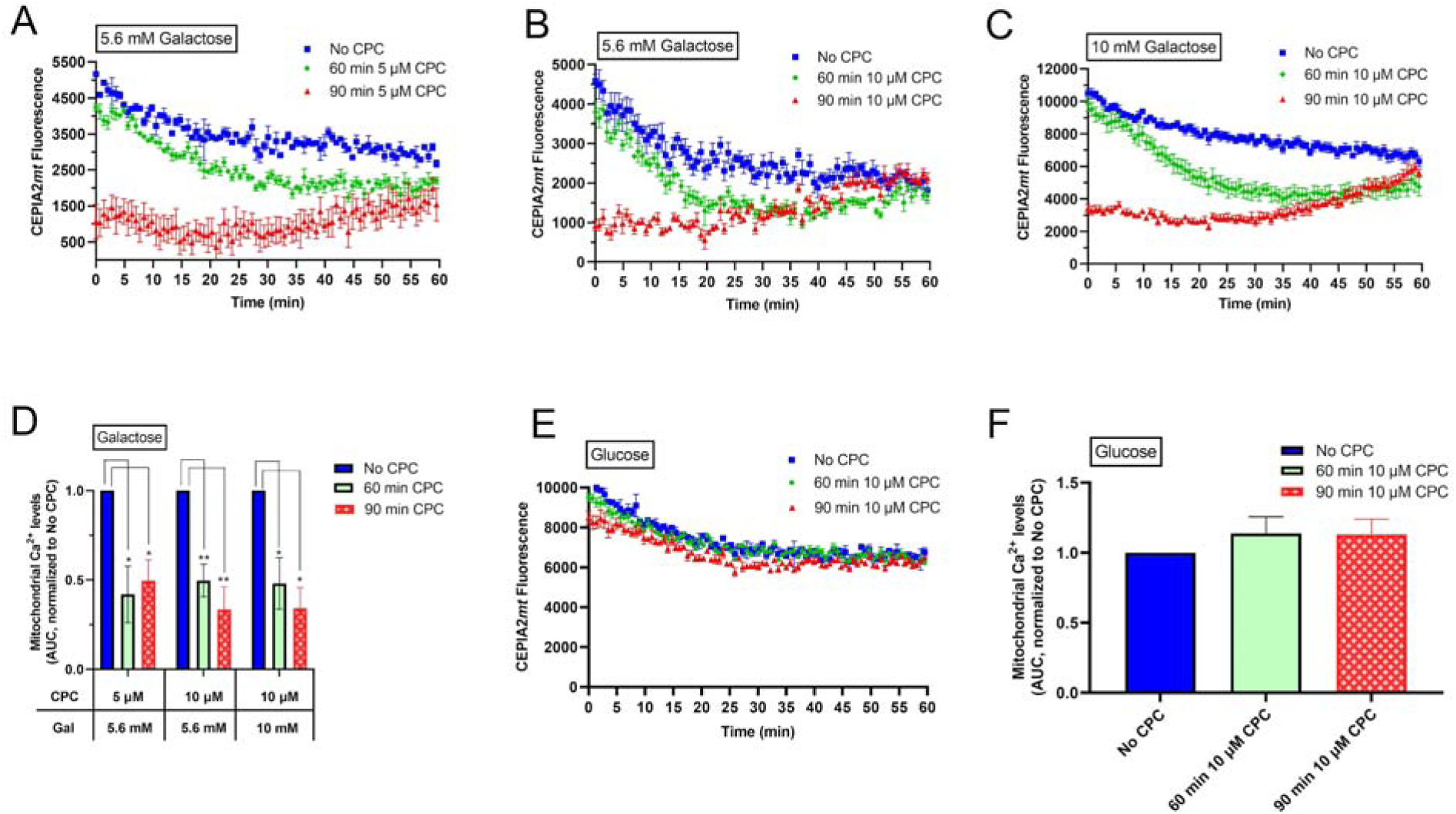
Effects of CPC on RBL-2H3 cell mitochondrial Ca^2+^ levels in galactose and glucose media. RBL cells were transfected with fluorescent CEPIA2mt construct and plated overnight. Cells were placed in galactose or glucose BSA-Tyrodes (BT) media and exposed to one of the conditions: “No CPC,” “60 min CPC,” “90 min CPC” as defined in methods. Representative CEPIA2mt fluorescence intensity curves of 5.6 mM galactose BT with 5 μM **(A)** and 5.6 mM galactose BT with 10 μM CPC **(B)** as well as 10 mM galactose BT with 10 μM CPC **(B).** Areas under the fluorescence curves (AUC) were determined and normalized to 0 µM CPC control for the various galactose (Gal) and CPC concentration combinations as indicated **(D)**. Representative CEPIA2mt fluorescence intensity curves of 5.6 mM glucose BT with 10 μM CPC **(E)**. AUC for the glucose BT group was determined and normalized to 0 μM CPC **(F)**. All values presented are mean ± SEM from at least three independent experiments, each with three replicates per CPC treatment. *p < 0.05, **p < 0.01, one-way ANOVA followed by Tukey’s post-hoc test.

In contrast, CPC addition to cells in glucose-containing buffer had no effect on mitochondrial Ca^2+^ levels, as indicated by a lack of change of CEPIA2mt reporter fluorescence (Figure 4E). AUC analysis of 10 μM CPC exposure (60 or 90 min) in glucose-containing buffer indicates no significant effects (Figure 4F).

### CPC disrupts mitochondrial morphology as measured with super-resolution microscopy

FPALM was utilized to precisely assess CPC effects on mitochondrial morphology at the nanoscale. A photoactivatable fluorescent marker of the outer mitochondrial membrane, Dendra2Tom20, was imaged in NIH-3T3 cells treated with 0 μM CPC (Control) or 5 μM CPC in galactose media. CPC induces a donut-like morphology in the mitochondria of NIH-3T3 cells (representative images in Figure 5). The presence of the donuts in the control and 5 μM CPC groups was quantified based on the number of observed donuts in each generated image (Figure 6).

**Figure 5.**
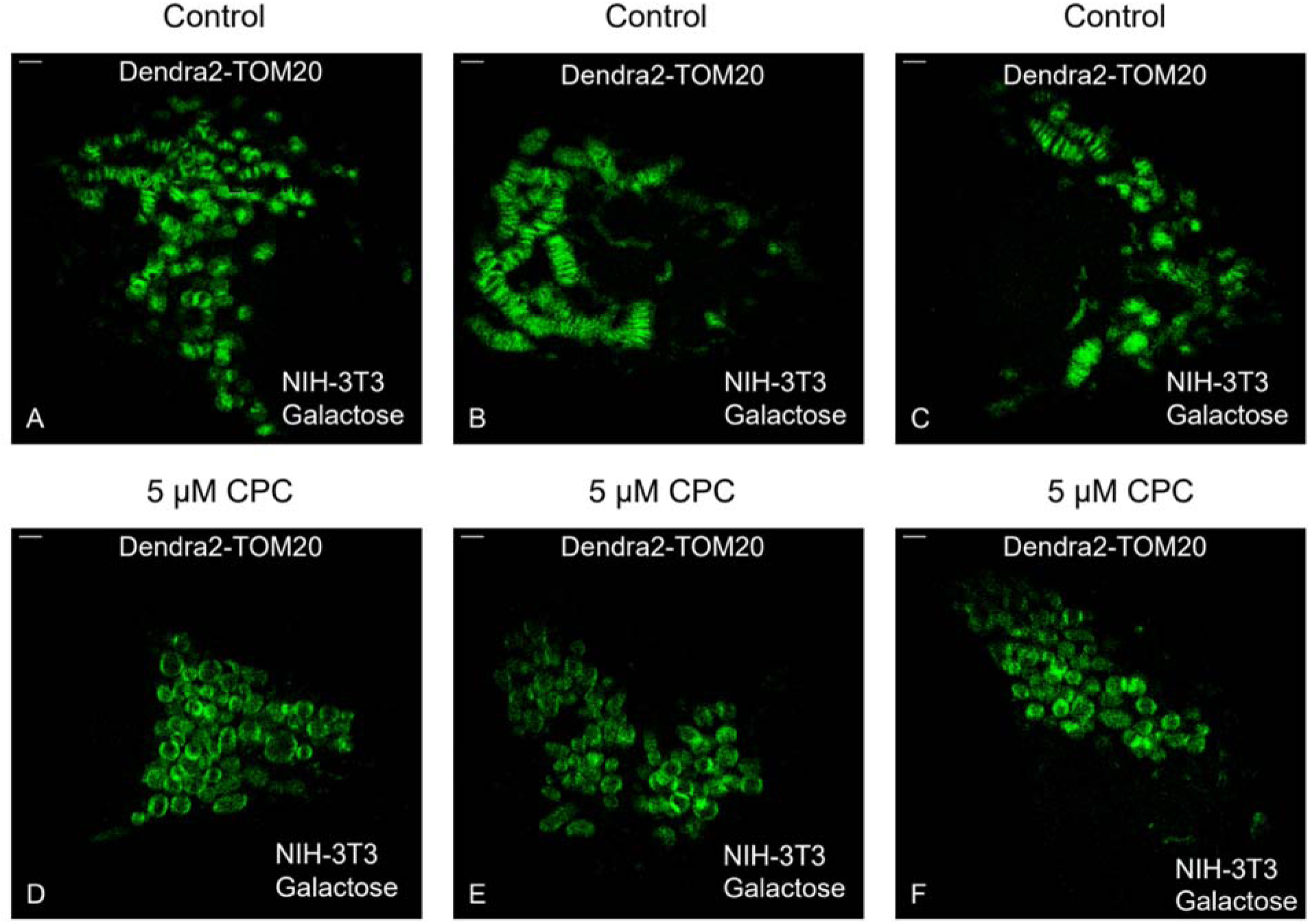
Representative super-resolution microscopy images of NIH-3T3 cell mitochondria structural effect following 0 μM CPC (Control) and 5 μM CPC exposure in galactose media for 60 min. FPALM images of live NIH-3T3 cell mitochondria expressing outer-membran marker Dendra2TOM20 (green) in Control 0 µM CPC **(A-C)** and 5 µM CPC **(D-F)** treated cells. The horizontal and vertical axes are defined as X (right), and Y (up), respectively. Shown are representative images from two days of imaging during which n = 30 of control cells and n = 30 of 5 μM CPC treated cells were imaged. Scale bars = 1 µm.

**Figure 6.**
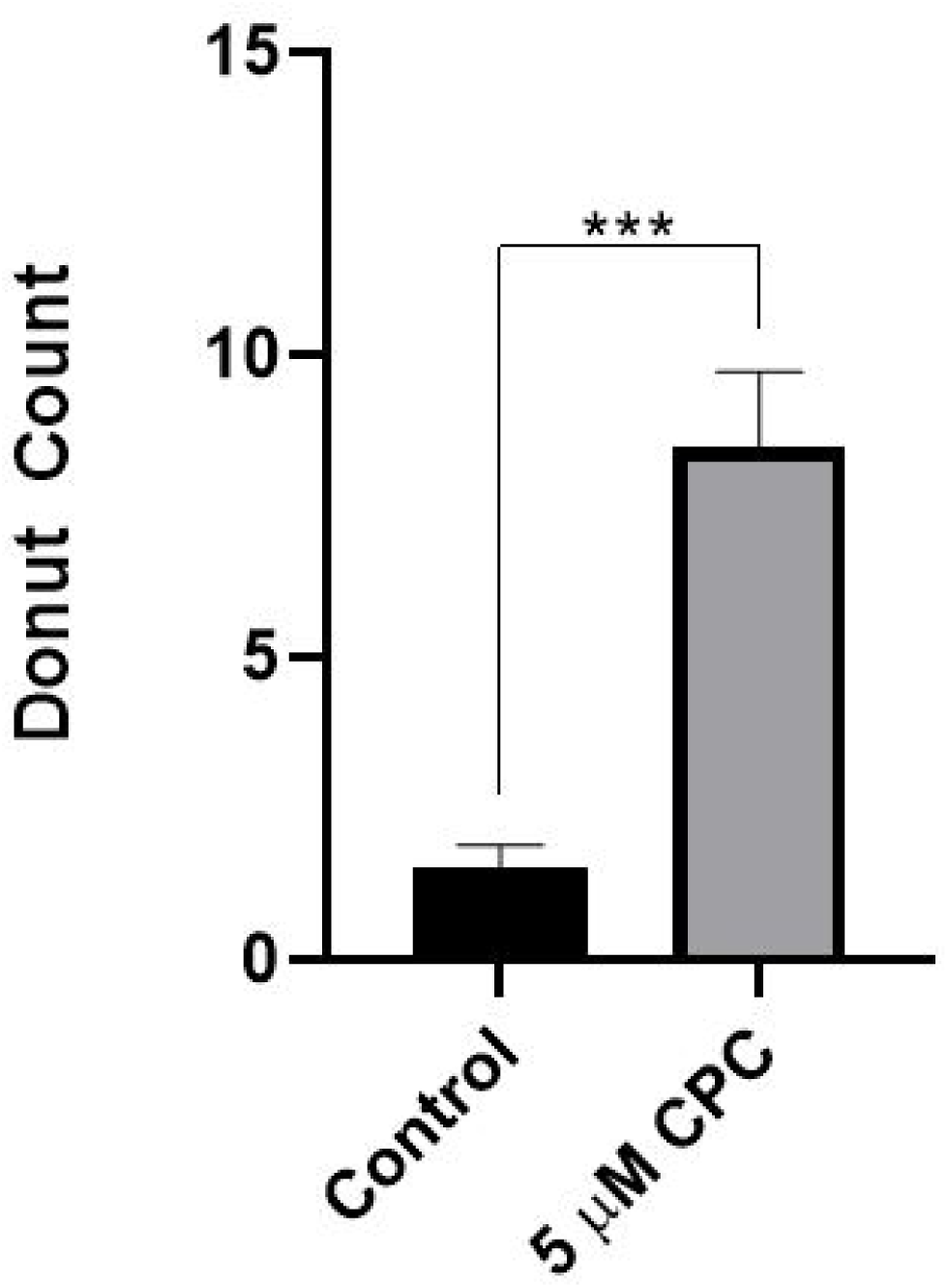
Mitochondrial donut formation in super-resolution microscopy images of NIH-3T3 cells following 0μM CPC (Control) and 5 μM CPC in galactose media exposure for 60 min. Live NIH-3T3 cells expressing mitochondria outer-membrane marker Dendra2TOM20 were imaged using FPALM. Donut shapes were observed in images recorded by three separate researchers via a blinded, randomized analysis in control (n = 25) and 5 μM CPC (n = 25). The average counts for each image were gathered and mapped back to the control or 5 μM groups. Significance is represented by ***p < 0.001 as determined by a Mann-Whitney test.

The morphological variation was quantified using the Fast Fourier Transforms of FPALM renders (Figure 7), where the control group can be observed to have a more symmetric distribution of spatial frequencies compared to the 5 μM CPC group. The donut shapes present in the sample had specific features and spatial frequencies typically within the range of 0.1–0.45 μm^-1^ (Figure 7). Furthermore, when donut-like shapes were observed, two of the sides of the donut (nominally defined as X; Figures 5 and 7) were typically observed to have brighter fluorescence than the other two sides (nominally defined as Y; Figures 5 and 7). Thus, the differences in the histogram of spatial frequencies as a function of X and Y were able to capture the differences in mitochondrial morphology (Figure 7C). The probability distributions for the dimensional ratios were statistically significant (p < 10^-10^) using a Kolmogorov-Smirnov test) comparing control and 5 μM CPC.

**Figure 7.**
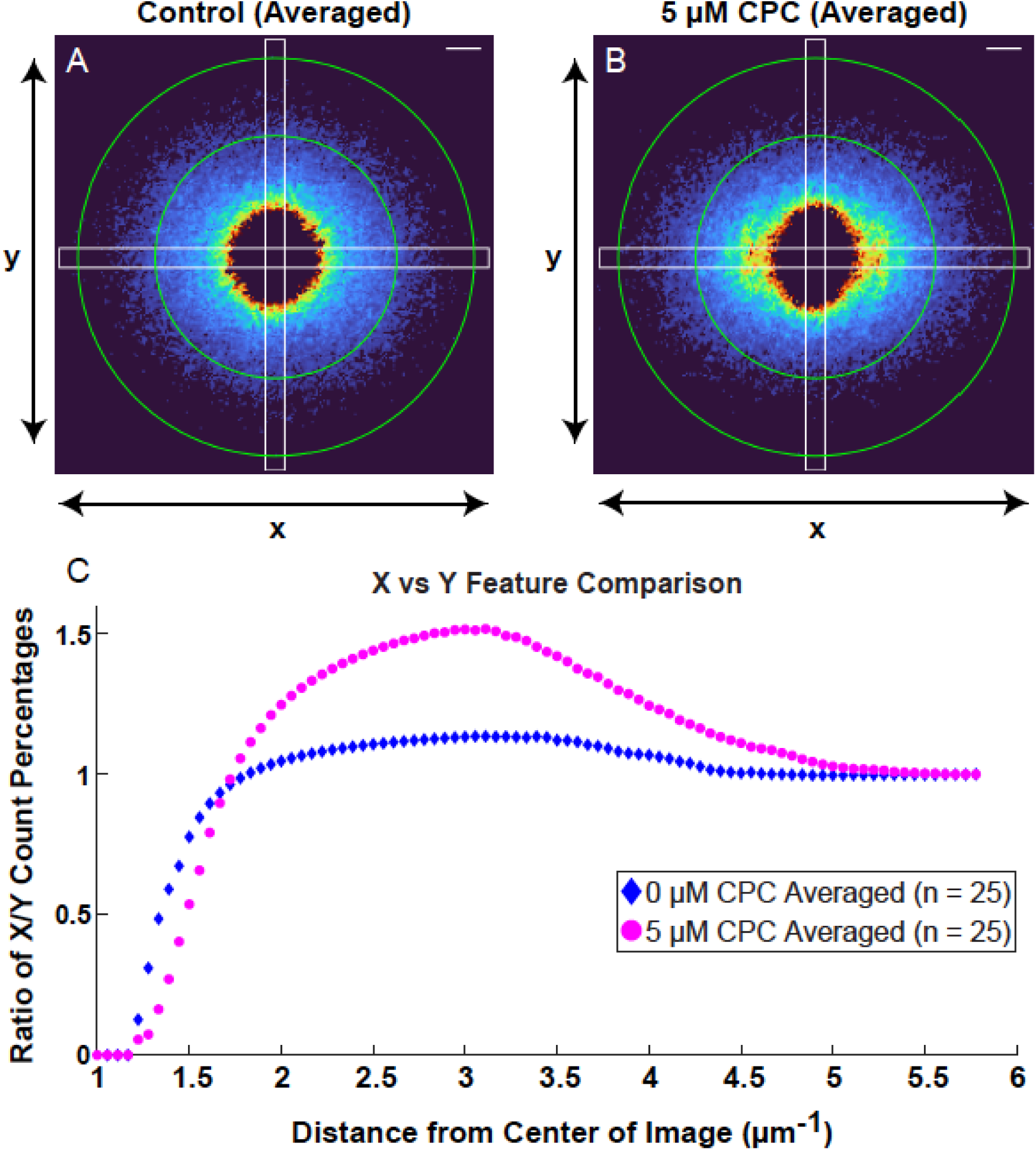
Fourier Transform Analysis of NIH-3T3 Mitochondrial Morphology following 0μM CPC (Control) and 5 μM CPC exposure in galactose media for 60 min. Averaged Fourier transforms of cellular images for control (n = 25) **(A)** and 5 μM CPC (n = 25) **(B)** samples after thresholding with a scale bar representing a wavenumber k of 1 μm^-1^. Images were of live NIH-3T3 cells transfected with Dendra2TOM20 and generated via FPALM rendering. Green circles in **(A-B)** represent an estimated region for features between 190-320 nm in size as the spacing for some of the most relevant structural features. This was used to estimate a lower threshold for features to be included, and an upper threshold was added to remove features at longer length scales represented at the center of the images. The white rectangles in **(A-B)** correspond to the regions selected for calculation in **(C)**. The graph in **(C)** plots the fraction of points within the rectangle spanning the X-direction contained within a certain distance from the center as a ratio with the same fraction in the Y-direction. This was used to quantify the elongation of the shape at different distances with a near circular shape corresponding to a ratio of 1. A Kolmogorov-Smirnov test returned a significance of p *<* 10^-10^ for the two curves and similarly for the two curves starting at a distance ≥ 2 μm^-1^ to assure enough data was collected before comparison.

## Discussion

In this study, we have revealed the mitochondrial toxicity of the common personal care and food chemical CPC towards multiple cell types, including primary human keratinocytes. We have shown that CPC inhibits mitochondrial ATP production (Figure 1), inhibits OCR (Figure 2), and causes efflux of mitochondrial Ca^2+^ (Figure 4). The mitochondrial toxicity of CPC seems to be more potent than its inhibition of mast cell signal transduction (Figure 3).

Using live-cell super-resolution microscopy, we show that CPC deforms mitochondria at the nanoscale, inducing a toroidal, “donut” morphology (Figures 5-7). The formation of “donuts” is a mode of fission of mitochondrial networks. Mitochondrial shape is key for healthy mitochondrial function and plays a role in disease development (Youle and Van Der Bliek, 2012). Abnormal mitochondrial structures including donuts are associated with disease etiology (Youle and Van Der Bliek, 2012), including insulin resistance (Jheng *et al*., 2012), bipolar disorder (Cataldo *et al*., 2010), memory problems in monkeys (Hara *et al*., 2014), Parkinson’s disease (Cui *et al*., 2010; Bhandari *et al*., 2014; Pozo Devoto and Falzone, 2017), and problems with embryonic development (Chen *et al*., 2003).

Donut patterns observed in 5 μM CPC samples tended to have an x- or y-directional bias at certain distances (Figure 7B). As the Fourier transforms are averaged over the set of cells (n = 25 per treatment type), it is assumed that directional bias from cell orientation will be averaged over as in the 0 μM control (Figure 7A). Membrane-associated fluorophores have often been observed to be preferentially excited by a given direction of laser polarization when their transition dipoles align with that laser polarization, leading to asymmetric intensity distributions in images of spherical structures (Krebs *et al*., 2004; Brasselet, 2011). Here, we expect that the membrane-associated TOM20-Dendra2 molecules are preferentially excited by a similar process, leading to the asymmetric fluorescence signals when spherical (donut-like) membranes are present (Figure 5). The Fourier transform of the images is then expected to be asymmetric when the donuts are present, and this is confirmed by the Fourier transforms of experimental images (Figure 7A and B). Hence, we quantify the asymmetry of the Fourier transform as a measure of the degree to which the donut shapes are present on the average, as a function of CPC exposure.

The mechanism underlying CPC disruption of mitochondrial nanostructure and formation of donuts (Figure 5) is unknown. Mitochondrial uncouplers including CCCP, FCCP, and TCS also cause donut formation, in multiple cell types (Liu and Hajnoczky, 2011; Ding *et al*., 2012; Giedt *et al*., 2012; Weatherly *et al*., 2018). However, because CPC possesses no ionizable proton and thus cannot act via an uncoupling process, its biochemical process of structural deformation is necessarily distinct from that of uncouplers. Because CPC may be an electron transport chain (ETC) inhibitor (Chávez and Concepcion, 1982; Hess *et al*., 2006; Datta *et al*., 2017b), CPC’s mechanism may be related to its ETC inhibition: antimycin A and rotenone also induce donut formation, potentially via mitochondrial reactive oxygen species (ROS) generation (Plecita-Hlavata *et al*., 2008; Giedt *et al*., 2012; Ahmad *et al*., 2013; Bulthuis *et al*., 2019). Another potential mechanistic explanation for CPC-induced donuts involves CPC (3 & 10 μM; 2 hr) activation of AMP-activated protein kinase (AMPK), which has been demonstrated in liver cells (Allen *et al*., 2020). Interestingly, inositol, which prevents aberrant AMPK activation, also prevents mitochondrial fission (Hsu *et al*., 2021). Because CPC both activates AMPK (Allen *et al*., 2020) and causes donut formation (Figure 5), AMPK stimulation may be involved in the mechanism underlying CPC formation of mitochondrial donuts.

To compare the mitochondrial toxicity of CPC to that of known mitotoxicants, in Table 1 we list EC_50_ values for inhibition of mitochondrial ATP production in RBL-2H3 cells, in galactose media, measured under nearly identical conditions. As a striking example, the canonical mitochondrial uncoupler CCCP exhibits an EC_50_ of 1.2 µM in this cell line in the same media type as the CPC experiments (with the only differences being media without BSA and with an incubation time of 2 hr in the CCCP experiments) (Weatherly *et al*., 2016), indicating that CPC (EC_50_ of 1.7 µM) is basically as mitotoxic as CCCP. Another canonical mitochondrial toxicant, 2,4-dinitrophenol (DNP) (Loomis and Lipmann, 1948), has an EC_50_ of ∼314 µM, which is ∼184-fold *less* potent as a mitotoxicant than CPC; DNP was a diet drug that worked via mitochondrial uncoupling and that was banned in 1938 in the USA due to its severe health effects (Grundlingh *et al*., 2011). Another antibacterial agent recently determined to be mitotoxic (Weatherly *et al*., 2016), TCS, has an EC_50_ of ∼8.6 µM, which is also *less* potent than CPC, by ∼5-fold.

**Table 1.**

| Mitotoxicant | EC<sub>50</sub> Values for Inhibition of Mitochondrial ATP Production |
| --- | --- |
| CCCP<sup>1</sup> | 0.8 – 1.6 μM (95% Confidence Interval) |
| CPC<sup>2</sup> | 1.5 – 2.0 μM (95% Confidence Interval) |
| TCS<sup>1</sup> | 7.5–9.7 μM (95% Confidence Interval) |
| DNP<sup>3</sup> | 389–677 μM (95% Confidence Interval) |

<sup>1</sup>Weatherly *et al.*, 2016
<sup>2</sup>Figure 1
<sup>3</sup>Weatherly *et al.*, 2018

CPC reduces O_2_ consumption rate (Figure 2). In RBL-2H3 cells, CPC at the ∼EC_50_ value for mitochondrial ATP suppression, 1.75 μM, suppressed OCR to 51% ± 8% of that of untreated cells (Figure 2B), supporting the concept that CPC’s suppression of ATP production as a direct consequence of its ETC inhibition. This magnitude of decrease is comparable to the amount of *increase* in OCR seen in RBL-2H3 cells exposed to canonical uncoupler CCCP (1 μM): 150% ± 10% (SEM) (Weatherly *et al*., 2016), although CPC depresses OCR while CCCP stimulates OCR due to differences in mechanisms of toxicity. Also, in primary human keratinocytes, CPC at the ∼EC_50_ value for mitochondrial ATP suppression, 1.25 μM, suppressed OCR to 32.1 % ± 0.1% of that of untreated cells (Figure 2D), lending further support to the concept of CPC’s suppression of ATP production as a direct consequence of inhibition of oxygen consumption (Figure 2D). This CPC mitochondrial toxicity of mitochondrial ATP and OCR inhibition may be universal, based on the 3 cell types (from different species and tissues) tested in this study plus those tested previously (Saladino *et al*., 1971; Datta *et al*., 2017b). If CPC suppresses mitochondrial ATP production in part via inhibition of mitochondrial complex I (Chávez and Concepcion, 1982; Datta *et al*., 2017b), its mitotoxicity suggests organismal toxicity, since Parkinson’s disease onset is linked with mitochondrial complex I inhibitors 1-methyl-4-phenyl- 1,2,3,6-tetrahydropyridine (Langston *et al*., 1983; Qureshi and Paudel, 2011) and rotenone (Sherer *et al*., 2003; Tanner *et al*., 2011). Further research into CPC’s biochemical mechanisms of OCR inhibition is needed to better characterize this chemical and inspire further animal model studies and potential epidemiological studies into human CPC exposure.

Beyond the proposed mitochondrial complex I inhibition, CPC lipid interference, specifically the documented CPC interference with phosphatidylinositol 4,5-bisphosphate, PIP_2_ (Raut *et al*., 2022), could also be implicated in mitochondrial toxicity. While mitochondrial lipid composition varies by cell type and by location (inner vs. outer membrane), phosphatidylinositols generally comprise a significant 5-20% of mitochondrial membrane composition (Daum, 1985; Stefanyk *et al*., 2010). PIP_2_ plays an essential structural role in mitochondria: its masking and inhibition of its signaling capabilities through overexpression of a PIP_2_-binding Pleckstrin homology domain caused increased levels of mitophagy and fission (Rosivatz and Woscholski, 2011). For instance, a study showed that relatively high doses of CPC destroy SARS-CoV2 viruses, the lipid envelopes of which are enriched in phosphatidylinositols (PI) (along with phosphatidylcholine and phosphatidylethanolamine), perhaps enriched in PIP_2_ (Saud *et al*., 2022). Thus, it is a plausible hypothesis that CPC destroys SARS-CoV2 by interfering in part with viral lipid envelope PIP_2_ (Saud *et al*., 2022) and that CPC could effect mitochondrial toxicity similarly by interfering with mitochondrial PIP_2_. Thus, the known interference of CPC with PIP_2_ (Raut *et al*., 2022) combined with PIP_2_’s structural role in mitochondria (Rosivatz and Woscholski, 2011) could be a mechanism underlying CPC’s mitochondrial toxicity and will be a topic of future study. We hypothesize that CPC mitotoxicity involves CPC interference with PIP_2_ from the mitochondrial membrane, causing mitochondrial fragmentation (Rosivatz and Woscholski, 2011).

An additional potential lipid-based mechanism underlying CPC mitotoxicity could involve PIP_2_. LDL receptor-related protein 1 (LRP1) recruits kinases that synthesize PIP_2_, which is enzymatically cleaved to produce diacylglycerol, needed to make cytidine-5’diphosphate-1,2- diacylglycerol (CDP-DAG) (Blunsom and Cockcroft, 2020), a substrate used to synthesize mitochondrial lipid cardiolipin (Chinnarasu *et al*., 2021). When LRP1 is knocked out in mice, mitochondria exhibit reductions in cardiolipin and PIP_2_, OCR inhibition, and mitochondrial structural defects (Chinnarasu *et al*., 2021). Thus, CPC interference with PIP_2_ may lead to reduced cleavage of it into DAG, leading to decreased production of CDP-DAG, and subsequent decreased levels of cardiolipin, which may reduce mitochondrial respiration. CPC interference with lipid synthesis may parallel the documented QAC inhibition of cholesterol synthesis (Herron *et al*., 2016).

The mechanism of CPC suppression of mitochondrial Ca^2+^ levels (Figure 4) is unknown. However, while mitochondrial proton ionophore uncouplers increase mitochondrial Ca^2+^ while concurrently suppressing ATP levels (and decreasing mitochondrial membrane potential [MMP]) (Weatherly *et al*., 2016; Weatherly *et al*., 2018), CPC exposure leads to *decreases* of both mitochondrial Ca^2+^ and ATP, indicating a mechanism of mitochondrial Ca^2+^ modulation distinct from that of uncouplers.

We found that CPC suppresses mitochondrial Ca^2+^ concentration only when mitochondrial ATP production was also inhibited (*i.e.*, in galactose media, not in glucose media, which allows glycolytic production of ATP; Figure 1). These data suggest that CPC causes mitochondrial Ca^2+^ efflux because it inhibits mitochondrial ATP production. One potential mechanism could be that CPC’s reduction of ATP levels results in malfunction of the mitochondrial Ca^2+^ uniporter (MCU), which works in energized cells to transport Ca^2+^ (when its cytosolic concentration is > 1 μM) into the mitochondrial matrix (Fan *et al*., 2020). For example, positively-charged ruthenium complexes inhibit MCU, albeit via an unknown mechanism (Ying *et al*., 1991; Woods *et al*., 2019). Another putative mechanism could be that CPC’s reduction of mitochondrial ATP production results in impairment of the ATP-dependent (Erecińska and Dagani, 1990) plasma membrane transporter Na^⁺^/K^⁺^-ATPase, which works in energized cells to maintain plasma membrane potential by catalyzing the exchange of three intracellular Na^+^ cations for two extracellular K^+^ cations (Nelson and Cox, 2017). Impairment of Na^⁺^/K^⁺^-ATPase would lead to cytosol buildup of Na^⁺^, then passive flow of Na^⁺^ down its concentration gradient through the cation-porous (Nelson and Cox, 2017) outer mitochondrial membrane, into the intermembrane space. At the inner mitochondrial membrane, the excess Na^⁺^ may then flow passively into the matrix via the ATP-*independent* Na^+^/Li^+^/Ca^2+^ exchanger (NCLX) (Palty *et al*., 2004). This inflow of Na^+^ could then cause mitochondrial Ca^2+^ efflux via NCLX (Palty *et al*., 2010), resulting in the decreased mitochondrial Ca^2+^ levels observed (Figure 4). Indeed, cytosolic Na^+^ increases can lead to efflux in matrix Ca^2+^ levels in cardiomyocytes (Liu and O’Rourke, 2008). NCLX function is reliant on MMP (Kostic *et al*., 2018). Such questions–CPC effects on MCU, NCLX, MMP, as well as on other Ca^2+^ transporters–will be topics of future research.

Apparent recovery of mitochondrial Ca^2+^ levels, for example in groups pre-exposed for 30 min to CPC, begins at ∼30 min of the plate reader time course, equating to a 60 min total of CPC exposure (Figure 4A-C). This refilling of mitochondrial Ca^2+^ occurs under conditions when mitochondrial ATP production is strongly inhibited (Figure 1). When mitochondrial ATP is not being produced, ATP synthase is not functioning, and thus H^+^ is not flowing back into the matrix through ATP synthase, and the matrix becomes relatively more alkaline (Nelson and Cox, 2017). In RBL-1 cells, MCU function is pH-dependent: MCU is activated at higher matrix pH levels (Moreau and Parekh, 2008). Thus, the observed mitochondrial Ca^2+^ recovery at later exposure times (Figure 4) may be due to activation of MCU.

Does CPC’s mitotoxic inhibition of ATP production (Figure 1) explain CPC’s inhibition of mast cell function (Raut *et al*., 2022)? We have found that mast cell degranulation is inhibited by CPC to the same degree, regardless of whether ATP is concurrently inhibited (in galactose media) or not inhibited (in glucose media) (Figure 3). Therefore, CPC’s ability to inhibit mast cell degranulation is independent of mitochondrial toxicity. This finding shows that CPC has additional cellular signal transduction targets of inhibition, to be probed in future work. The results in Figure 3 also demonstrate that CPC’s mitochondrial toxicity exceeds the potency of CPC’s immune cell signaling interference: the EC_50_ is ∼10 μM CPC for inhibition of degranulation (Figure 3) whereas the EC_50_ is 1.7 μM for CPC’s inhibition of mitochondrial ATP production (Figure 1), a value ∼6-fold lower under the same exposure conditions. Thus, CPC appears to be ∼6-fold more potently toxic to mitochondria than to general cellular signal transduction. However, if CPC inhibits ATP production through mitochondrial complex I inhibition (Datta *et al*., 2017b), this process could mask increased degranulation inhibition: mitochondrial complex I inhibition can lead to increased levels of ROS (Fato *et al*., 2009), which are known stimulators of mast cell degranulation (Swindle *et al*., 2004). Future research will examine CPC effects on various types of ROS, in mitochondria and elsewhere, to investigate this mechanistic avenue. Furthermore, the antiestrogenic effect of CPC after 24 hr exposure could interfere with the known stimulation of degranulation through estrogen exposure (Zaitsu *et al*., 2007), implicating a potential connection between estrogenic and immunogenic signaling inhibition. In the timeframe of a 90 minute exposure to CPC, other mechanisms are likely at play to cause levels of ATP lowered by ∼90% at 10 μM (Figure 1) without cytotoxicity.

In conclusion, we have shown that CPC acts as a mitotoxicant in multiple types of living cells including primary human cells, at concentrations ∼3,000-fold lower than those used in consumer products. CPC causes inhibition of mitochondrial ATP production, oxygen consumption, mitochondrial Ca^2+^ buffering (as revealed via a novel experimental method), and disruption of mitochondrial nanostructural integrity, as shown via FPALM super-resolution microscopy. This study is one of the first uses of super-resolution microscopy in the field of toxicology. These data provide insight into mechanisms underlying CPC disruption of mitochondrial function at doses relevant to human exposure via consumer and food products. In summary, we show that CPC is a mitotoxicant in mammalian cells (rat, mouse, and human), similarly potent as canonical mitotoxicant CCCP and more potent than other previously determined mitotoxicants, DNP and TCS. Likely due to its continual use since the 1930s (ACS, 2021), modern toxicology studies were never conducted to evaluate potential risks of CPC despite its continued existence in many consumer/cleaning/food uses. Its continued widespread use raises the importance of identifying any potential harmful effects. Our findings provide considerations for this chemical so that decision-makers (government regulators, medical authorities, consumers, companies, etc.) can make informed choices to weigh the risks and benefits of CPC use.

## Acknowledgments

We thank Roger Sher for the Dendra2-TOM20 construct, Komala Shivanna for providing access to shared lab materials and resources, David Winski for insightful discussions, Matthew Parent for MATLAB scripts, Patrick Fleming and Marissa Paine for help in ordering materials and lab support, Patricia Byard and Timothy Campbell for administrative assistance, Dr. Melody Neely and Caitlin Wiafe-Kwakye for allowing our lab to use their nanodrop, as well as Dr. Melissa Maginnis and Avery Bond for providing use of equipment.

This project was funded primarily by the University of Maine System Research Reinvestment Fund Grant Program Track 1 Rural Health and Wellbeing Grand Challenge (PI: Gosse), by a UMaine Medicine Seed Grant (PI: Gosse), and from a Bioscience Association of Maine (BioME) Seed Grant. This research was also supported the National Institutes of Health: National Institute of General Medical Sciences award numbers 1R15GM139070 (PI: Hess) and P20GM103423 (an Institutional Development Award). The Maine Technological Asset Fund (MTAF 1106 and 2061, PI: Hess) provided equipment. University of Maine student funding that supported this work includes a Graduate Student Government Grant, the Center for Undergraduate Research (CUGR), the Charlie Slavin Research fund of The Honors College.

## Abbreviations

Ag: antigen
AMPK: AMP-activated protein kinase
AUC: area under the curve
BSA: bovine serum albumin
BT: bovine serum albumin in Tyrode’s buffer
CCCP: carbonyl cyanide 3-chlorophenylhydrazone
CCW: cell culture water
CDP-DAG: cytidine-5’diphosphate-1,2-diacylglycerol
CI: confidence interval
CPC: cetylpyridinium chloride
DAG: diacylglycerol
EC_50_: Half maximal effective concentration
ETC: electron transport chain
FFT: Fast Fourier Transform
IgE: immunoglobin E
LHON: Leber’s hereditary optic neuropathy
MCU: mitochondrial Ca^2+^ uniporter
MMP: mitochondrial membrane potential
PIP_2_: phosphatidylinositol 4,5-bisphosphate
QAC: quaternary ammonium compound
RBL-2H3: rat basophilic leukemia
RFU: raw fluorescence unit
ROS: reactive oxygen species
SARS-CoV2: severe acute respiratory syndrome coronavirus 2
TCA: tricarboxylic acid cycle

