## Supplement for "Mechanisms of Antimicrobial Agent Cetylpyridinium Chloride Mitochondrial Toxicity in Rodent and Primary Human Cells: Super-resolution Microscopy Reveals Nanostructural Disruption"

**Table of Contents**

Effect of CPC on Mitochondrial ToxGlo Assay fluorogenic protease and luminescent ATP reactions in RBL-2H3 cells……………………………………………………………………2-3

Effect of CPC on Mitochondrial ToxGlo Assay fluorescence and luminescence readings in assay buffer……….……………………………………………………………………………………...4

Effect of CPC on Oxygen Consumption Rate Assay background phosphorescence in RBL-2H3 cells….…………………………………………………….………………………………………5

Effect of CPC on Oxygen Consumption Rate probe phosphorescence in assay buffer……….….6

Effect of CPC on Oxygen Consumption Rate probe phosphorescence, with or without cells, at 30 minutes…….……………………………….……………………………………………………...7

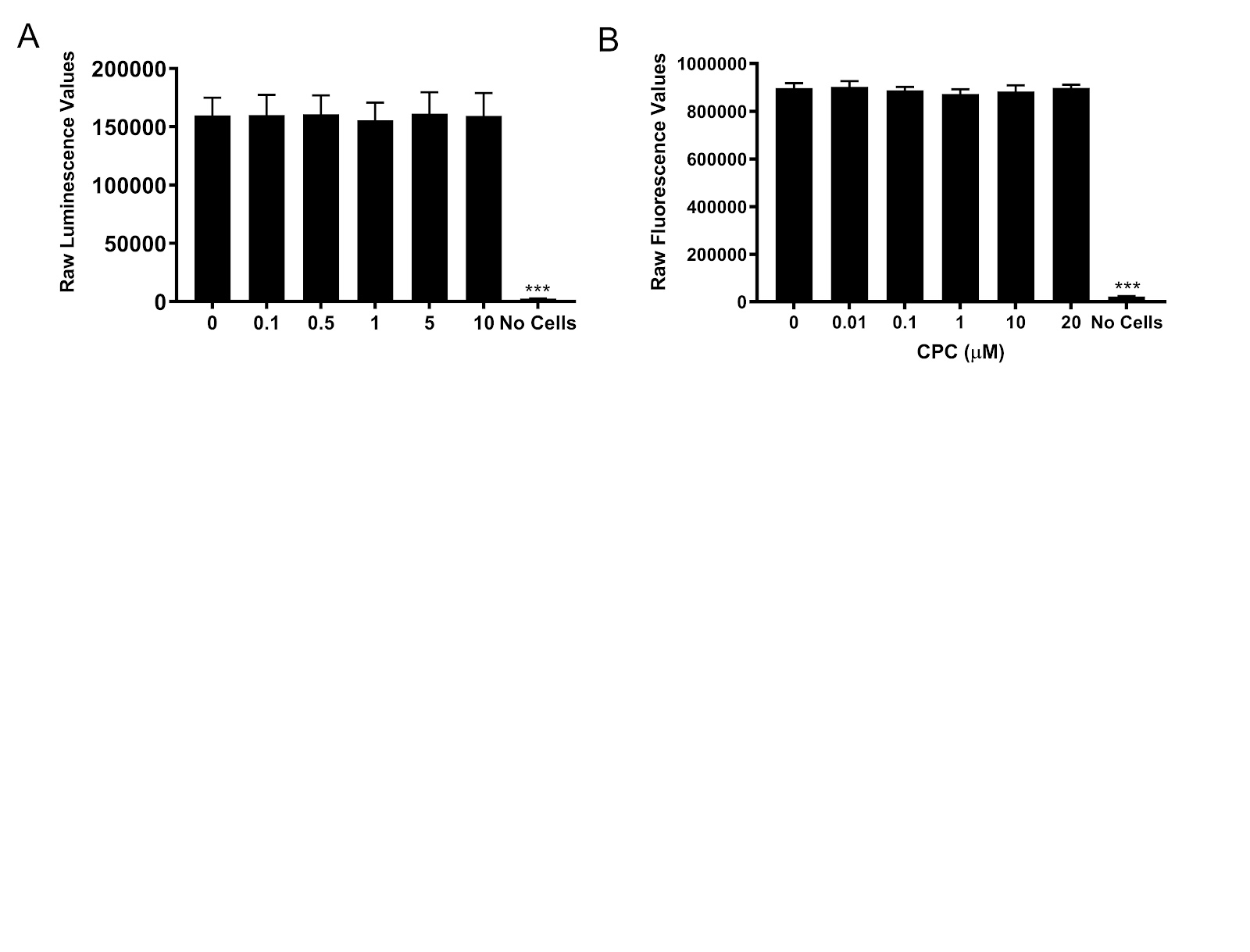

**Figure S1. Effect of CPC on Mitochondrial ToxGlo Assay fluorogenic protease and luminescent ATP reactions in RBL-2H3 cells.** To investigate the potential for CPC to interfere with these assays, ATP production and cytotoxicity were assessed under conditions which prevent the progression of cellular processes. Mitochondrial ToxGlo assay was performed as a control to determine whether CPC interferes with the reactions that form the luminescence from the ATP reagent **(A)** or the fluorescence from the cytotoxicity assay **(B)**.

**(A).** This experiment was done to assess whether CPC interferes with the luminescence detection of ATP in cell lysates, in contrast with typical ToxGlo experiments which test CPC effects on ATP production in living cells. For the ATP production assay control, luminescence was measured in cells lysed by ATP reagent prior to CPC-exposure. For this control experiment, an incubation of RBL-2H3 cells with media alone (No CPC) was done in place of the 60 min incubation with CPC. Next, ATP reagent was added to lyse the cells and to detect ATP, followed immediately by various concentrations of CPC, concomitant with the cytotoxicity reagent to simulate the conditions of the Mitochondrial ToxGlo Assay. The remaining luminescence measurement process was followed as in Methods.

**(B).** This experiment was done to assess whether CPC interferes with the cytotoxicity detection process itself, in contrast with typical ToxGlo experiments which test CPC effects on cytotoxicity in living cells. The cytotoxicity control experiment was conducted by lysing cells with digitonin prior to CPC-exposure. For the fluorescence control, all normal ToxGlo experimental steps were followed except cells were lysed with digitonin *prior to* CPC addition for a 60 min exposure.

Data shown as mean ± SEM of three independent experiments with triplicates in each experiment. No significance between any groups when compared to 0 µM CPC control, as determined by one-way ANOVA followed by Tukey’s post-hoc test.

**Results:** We tested whether CPC affects the ATP luminescence reaction in lysed cells, in an experiment in which CPC is not added until the cells are lysed **(A)**; these data show that CPC does not interfere with the detection of ATP. Additionally, we confirmed that CPC does not hinder the fluorescence reaction for cytotoxicity detection: CPC does not interfere with the protease–substrate reaction in this kit **(B)**.

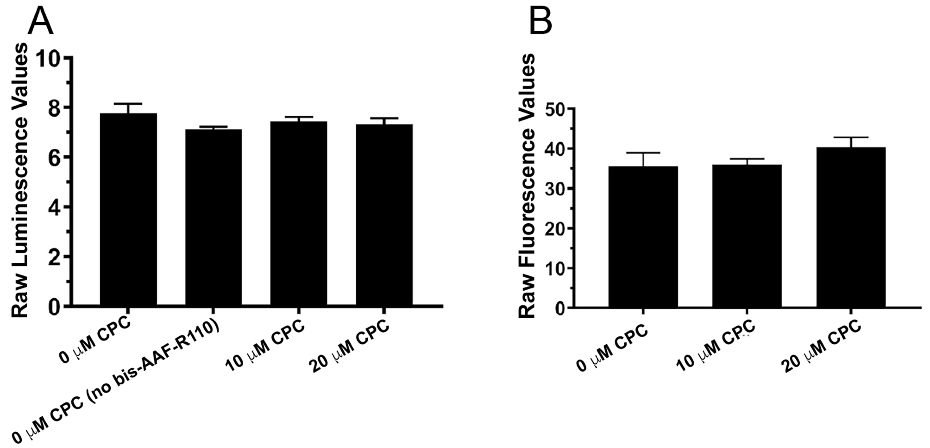

**Figure S2. Effect of CPC on Mitochondrial ToxGlo Assay fluorescence and luminescence readings in assay buffer.** No-cell controls were conducted to determine the effect of CPC on Mitochondrial ToxGlo Assay background luminescence **(A)** and fluorescence **(B)** readings. The Mitochondrial ToxGlo Assay was performed without the plating of cells **(A-B)**. All steps were performed as in Methods except for one sample with assay buffer without the addition of the bis-AAF-R110 cytotoxicity reagent. Data shown as mean ± SEM of three independent experiments with triplicates in each experiment. No significance between experimental groups when compared to 0 µM CPC control, as determined by one-way ANOVA followed by Tukey’s post-hoc test. These results indicate that CPC does not interfere with the processes of the ToxGlo assay.

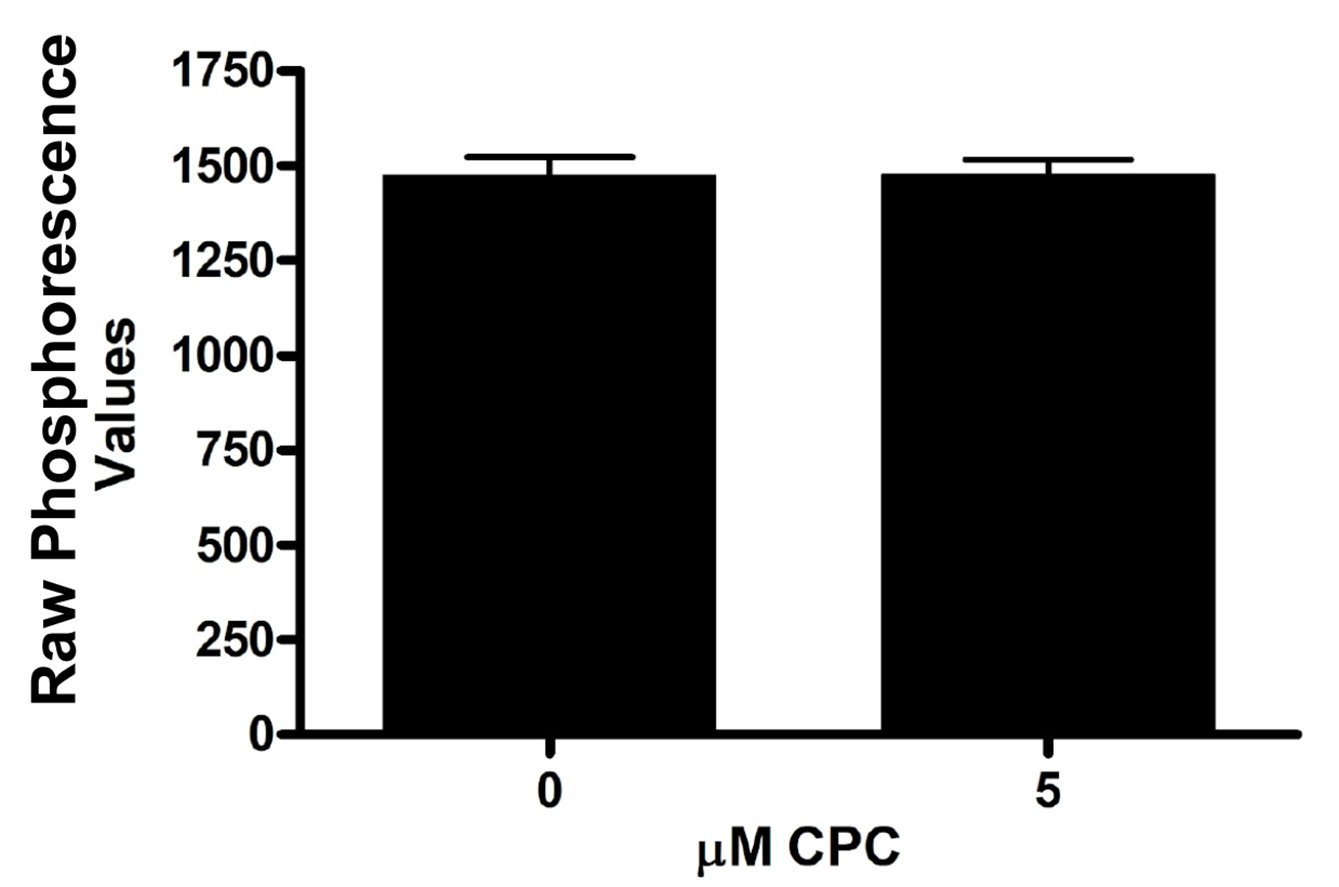

**Figure S3. Effect of CPC on Oxygen Consumption Rate Assay background phosphorescence in RBL-2H3 cells**. It is necessary to assess whether CPC interferes with the signal in RBL-2H3 cells from the phosphorescent probe used to measure OCR and to determine the integrity of the experiment. In order to do so, we first needed to evaluate whether CPC interferes with the background signal at the excitation and emission wavelengths utilized in this assay. Thus, a comparison was made of phosphorescence values averaged across all time points (0-180 min) for the wells containing cells but without OCR probe, ± CPC. This comparison would reveal whether the presence of CPC changes the background phosphorescence in the presence of cells and media but in the absence of probe. Data were produced from three independent experiments in quadruplicate and averaged across each individual time point. No significance between groups was detected via one-sample t-test. Thus, CPC does not alter the background (no-probe) phosphorescence of samples with cells in the OCR readings.

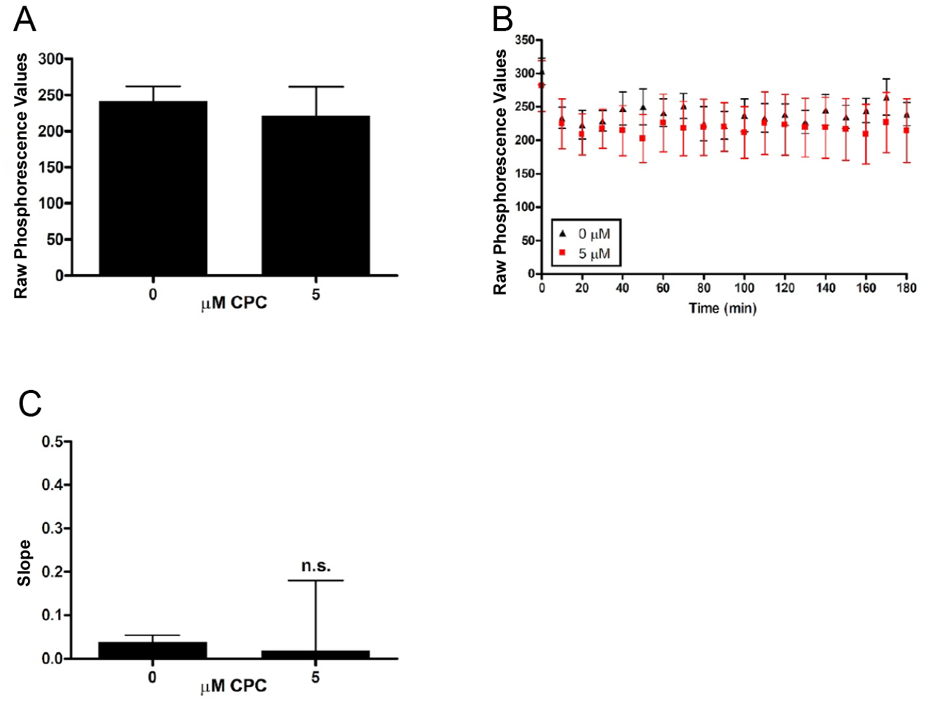
**Figure S4. Effect of CPC on Oxygen Consumption Rate probe phosphorescence in assay buffer.** To assess whether CPC affects OCR probe phosphorescence in the absence of cells, experiments with probe but without cells were conducted. Background phosphorescence (without cells, without probe) in presence of 0 μM and 5 μM CPC was averaged across the 180 min and subtracted from the respective samples that did contain probe. Data from two independent experiments, each experimental group run in duplicate **(A-C)**. The phosphorescence averaged over time for the wells containing probe, thus background-subtracted, are shown as mean ± SEM **(A)**. These same data were also plotted as a time course, mean ± SEM **(B)**. The slopes of the lines in **(B)** were analyzed from time point 30-180 min and plotted as a bar graph **(C)**. Data are not significant, as determined by one-sample t-test **(A, C)**. Thus, CPC does not change the OCR probe phosphorescence of samples without cells, showing that the OCR cellular data (Figure 2) is truly due to CPC effects on cellular oxygen consumption rates.

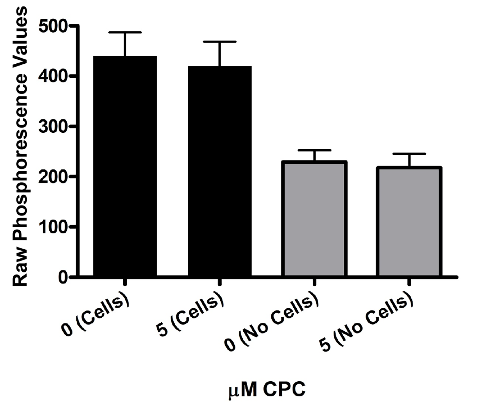

**Figure S5. Effect of CPC on Oxygen Consumption Rate probe phosphorescence, with or without cells, at 30 minutes.** The phosphorescence values of the OCR probe, for experiments both with and without cells, at an early time point t = 30 min were plotted. This time point was chosen because it is the starting point for the OCR experiments and because any CPC interference would be detectable by 30 min incubation. Experiments with cells were conducted on three experimental days (with four replicates per sample per day) whereas experiments without cells were performed on two days (with duplicates per sample per day). Data are not significantly different, as assessed via one-way ANOVA followed by Tukey’s post-hoc test. This result indicates that CPC does not interfere with the phosphorescent probe signal at the start of the OCR experiments, with or without cells.
